# On the Curvature and Relaxation of Microtubule Plus-end Tips

**DOI:** 10.1101/2025.05.23.655844

**Authors:** Tomasz Skóra, Jiangbo Wu, Daniel Beckett, Weizhi Xue, Gregory A. Voth, Tamara C. Bidone

## Abstract

Microtubules are essential cytoskeletal components with a broad range of functions in which the structure and dynamics of their plus-end tips play critical roles. Existing mechanistic models explain the tips curving dynamics in different ways: the allosteric model suggests that GTP hydrolysis induces conformational changes in tubulin subunits that destabilize the lattice, leading to protofilament curving and depolymerization, while the lattice model posits that GTP hydrolysis directly destabilizes the microtubule lattice. However, the effect of GTP hydrolysis on the curving dynamics of microtubule tips remains incompletely understood. In this study, we employed a multiscale modeling approach, combining all-atom molecular dynamics simulations with Brownian dynamics simulations, to investigate the relaxation of microtubule plus-end tips into curved configurations. Our results show that both GDP- and GTP-bound tips exhibit an outward bending of protofilaments into curved, ram’s horn-like structures, characterized by a linear relationship between curvature and distance from the plus-end tip. These observations align with experimental cryo-ET images of microtubule plus-end tips in different nucleotide states. Collectively, our findings suggest that the outward bending of protofilaments at the plus-end tip is an intrinsic feature of microtubules, independent of the nucleotide state.

**SIGNIFICANCE:** Understanding how microtubules change shape is crucial for elucidating key cellular processes such as cell division and shape maintenance, which are fundamental to both physiological function and disease progression. This study supports the concept that the microtubule plus-end tip relaxation does not align with models that couple shape changes to GTP hydrolysis, at least for the topmost tubulin heterodimers. By interfacing bottom-up multiscale modeling — using the longest reported atomistic molecular dynamics simulations of microtubule tips— with existing cryo-ET data, it is shown that protofilament bending operates independently of nucleotide hydrolysis and likely depolymerization. These findings highlight the need for a new conceptual framework that separates GTP hydrolysis from microtubule tip flaring.

## 1. INTRODUCTION

Microtubules (MTs) are essential cytoskeletal components with a wide range of biological functions, including providing structural support to cells (1), facilitating intracellular transport (2), enabling chromosome segregation during cell division (3), and aiding motility in structures like cilia and flagella (4). Their versatility in supporting these different processes across nearly all eukaryotic cells is attributed to their dynamic nature, including changes in shape. The MT plus-end tip regulates actin filaments polymerization during mitosis (5), contributes to the turnover of focal adhesions, to the establishment of the direction for cell migration (6–9), and exhibits dynamic instability — a stochastic switching between periods of growth, pause, and shrinkage (10, 11). During these processes, MTs’ plus- end tips can change shape, bind different intracellular targets including end binding proteins (12), generate mechanical force (1) and couple with the actin filaments (13). Understanding how the MT tip undergoes structural relaxation is important for elucidating fundamental biophysical mechanisms of cells.

MTs are composed of α-β tubulin heterodimers arranged in a hollow cylinder. These dimers form protofilaments (PFs) that align parallelly with screw symmetry, resulting in the cylindrical MT lattice. During MT assembly, the plus-end tip elongates through incorporation of GTP-tubulin dimers.

Subsequently, the β-tubulin-bound GTP are hydrolyzed to GDP (14). MTs with GDP-bound β-tubulins are less stable and disassemble while those presenting a stabilizing GTP-cap at the plus-end continue to grow (15–19). The rate of GTP hydrolysis is dependent upon interdimer lattice compaction related to how close a given dimer is to the plus-end tip – less compacted lattices at the tip hydrolyze GTP less quickly than more compacted lattices in the “bulk” of the MT (20). The instability of GDP MT lattices is often associated with the relaxation of PFs’ intrinsic curvature leading to lattice expansion, and outward- bending of protofilaments with formation of ram’s horn-like structures (21, 22). As a result, the two processes — depolymerization and PF curvature relaxation — have been generally considered related.

There are two prevailing models that explain the relationship between depolymerization and PF curvature relaxation at GDP-MT tips: the allosteric model and the lattice model. The allosteric model posits that GDP-tubulin is more bent than GTP-tubulin, creating strain in the MT lattice that leads to destabilization (22–24). In contrast, the lattice model attributes the instability of GDP-PFs and their greater bending compared to GTP-PFs to weaker lateral interactions among tubulin monomers in the lattice (25, 26).

Recent research challenges the conventional view that MT plus-end tip relaxation is directly linked to depolymerization, suggesting instead that these processes should be considered independent mechanisms. Evidence supporting this revised perspective includes several key observations. First, in both *in vitro* and *in vivo* studies, MT plus-end tips show outward bending during both depolymerization and polymerization, regardless of the nucleotide state (19, 27–34). This indicates that tip relaxation is not exclusive to depolymerization events. Second, GDP- and GTP-tubulins exhibit similar curvature, suggesting that nucleotide state does not determine tip shape (35–38). Finally, tubulin ancestors such as FtsZ proteins, which lack the MT lattice structure, display similar dynamic instability, indicating that dissociation from the lattice is not a hallmark of MT depolymerization (39–41). Together, these findings support a revised model in which tip relaxation is a distinct structural process that can occur independently of depolymerization, prompting a reevaluation of how mechanical transitions at the MT tip are understood.

Capturing the relaxation of a single MT tip experimentally is challenging due to the static nature of these structures in imaging, and uncertainties introduced by rapid freezing techniques and the low signal-to- noise ratio caused by averaging effects. All-atom Molecular Dynamics (MD) simulations are also limited in studying MT relaxation due to the large timescales, which extend beyond microseconds (42).

However, without detailed atomistic information, understanding the nature of MT plus-end tip relaxation remains difficult. Recent modeling efforts have combined all-atom and coarse-grained models to study the interplay between radial (19) and out-of-plane relaxation modes (38, 42, 43) of the MT PFs (34), and analyzed the emergence of bent PF clusters and sheets (34, 42–46). Despite these advancements, current models still struggle to capture the linear dependence of bending angle on distance from the tip (19).

Building on these limitations, we developed a multiscale modeling approach that combines all-atom MD with bottom-up coarse-graining (CG-ing), allowing us to capture long-time dynamics while retaining detailed atomistic information. To validate our model, we compared the sampled morphologies with cryogenic electron tomography (cryo-ET) images. Our results confirm that MT tips in both nucleotide states exhibit similar curved morphologies and bending behaviors, including protofilament clustering and partial out-of-plane curving, while also capturing the linear dependence of curvature on distance from the tip. By learning interaction potentials from all-atom simulations and without assuming differences in curvature or lateral interactions between dimers in different nucleotide states, our multiscale model highlights consistent features of MT tip relaxation across different nucleotide states, advancing the multiscale modeling of MT behavior from the molecular to the macromolecular level.

## 2. METHODS

### 2.1. Molecular dynamics simulations

We constructed initial configurations for the all-atom MD simulations based on the cryo-EM crystal structures of MT patches comprising 3 2-heterodimer-long PFs in GDP-bound (PDB 6DPV) and GMPCPP-bound (guanylyl-(α-β)-methylene-diphosphonate) (PDB 6DPU) states (47). Missing residues were added and their positions were optimized using MODELLER (48). The GMPCPP, a nonhydrolyzable analog of GTP, was converted to GTP by exchanging the methylene (CH_2_) group between the α- and β-phosphate with an oxygen atom.

The 8-heterodimer-long, 14-PF MT plus-end tip structures were generated from the patches leveraging MT’s screw symmetry, using a procedure based on the earlier work by Tong and Voth (49). Firstly, the patches were extended longitudinally one heterodimer at a time using the alignment feature of VMD (50): bottom dimers of one patch were aligned with the top dimers of the other patch. Secondly, these elongated patches were extended laterally one PF at a time to capture MT’s lateral curvature: the leftmost PFs of one patch were superimposed with the rightmost ones from the other patch. Finally, a seam region formed naturally between the 1st and the 14th PF, exhibiting a longitudinal pitch of three tubulin monomers.

We applied periodic boundary conditions, with a simulation box of size 495 Å × 495 Å × 822/842 Å for MTs in GDP/GTP states, respectively. We aligned the major MT axis with the z axis of the Cartesian coordinate system. Such box dimensions left 20 Å of space for PFs’ bending. Water was included explicitly using the TIP3P model with 150 mM KCl. The GDP-MT system contained 20,483,435 atoms comprising 1,513,680 protein atoms, 6,309,337 water molecules and 41,744 ions. The GTP-MT system contained 20,981,887 atoms comprising 1,514,240 protein atoms, 6,475,037 water molecules and 42,536 ions.

We used GROMACS version 2019.6 (51) for performing the energy minimization, equilibration and the production runs of the all-atom MD simulations. We employed the CHARMM36m force field with CMAP correction (52). We accounted for long-range electrostatic interactions with Particle Mesh Ewald method (53). We minimized the energy using the gradient descent algorithm in two stages, with a maximum displacement of 0.01 (first stage) and 0.2 Å (second stage). Equilibration was performed at 1 atm and 310 K in the NVT ensemble for 100 ps and subsequently in NPT ensemble for 500 ps, upon gradual removal of the position restraints from 10 to 2 kJ/mol/Å^2^. We immobilized heavy atoms of 14 bottommost α-tubulins in the production NPT runs, assuming that our system is an extreme tip of much longer MT. We used leap-frog integrator and controlled the pressure by using the Parrinello-Rahman barostat (54) and the temperature using canonical velocity rescaling approach (55).

The production runs were performed for 4 μs with a timestep of 2 fs. During the runs, simulation box size was increased to fit the increasingly curved and lengthening tips, and water and ions were added to the system to maintain its ionic strength (Fig. S1 and Tab. S1). We applied the same initial energy minimization and equilibrium protocol to the system at each box resizing before continuing the run.

### 2.2. Coarse-grained mapping

To simulate timescales beyond 4 μs at low computational cost, we mapped the all-atom structure of the MT into a CG representation. A CG mapping provides the rules for projecting an all-atom structure into its corresponding CG counterpart (56, 57). Using the OpenMSCG (58) cgmap utility, to each tubulin monomer we attributed a single bead located at its center of geometry. The resultant CG representation includes a total of 224 beads, half of them representing α-tubulin, and the other half – β-tubulin.

Following the all-atom structure, the beads are organized in 14 16-tubulin-long PFs

### 2.3. Brownian dynamics simulations

To evaluate MT tip flaring at timescales beyond 4 µs, we simulated time evolution of the MT model using Brownian dynamics (BD) (59–61). We propagated BD with the first-order forward Euler scheme:

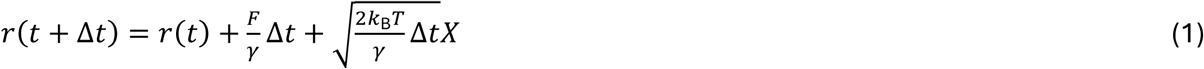

where *r* is position, *t* is time, *Δt* is timestep, *k*_B_ is Boltzmann constant, *T* is temperature, *γ* is friction coefficient, *F* is deterministic force, and *X* is standard normal random variable.

BD is rooted in Langevin dynamics, however, it assumes its overdamped regime, i.e., complete momentum relaxation during a single propagation step due to the fluid viscous drag. The large timestep *Δt* = 0.1 ps allows to ignore the dynamic memory kernel (62). In simulation starting from MT tubular form, we used *Δt* = 0.01 ps.

To perform BD simulations, we used LAMMPS (63) software. For computational efficiency, we neglected hydrodynamic interactions (64).

### 2.4. Generalized Rotne-Prager-Yamakawa approximation

Applying the Stokes law, we obtain the effective translational friction coefficient:

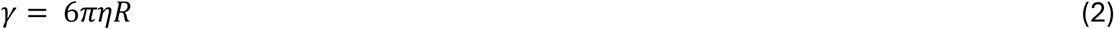

where *η* is fluid viscosity, and *R* is hydrodynamic radius, which depends on a particle’s size and shape.

In generalized Rotne-Prager-Yamakawa (GRPY) approximation, a macromolecule is represented as a conglomerate of beads of arbitrary radii, and treated as a rigid body (65). Firstly, the N-body friction matrix of such N-bead conglomerate is calculated (65). Secondly, it is projected onto the 3-dimensional subspace using the rigid body constraints. Finally, the friction matrix is partially inverted, yielding the rigid body grand mobility matrix. The hydrodynamic radius is given by its partial trace, i.e.:

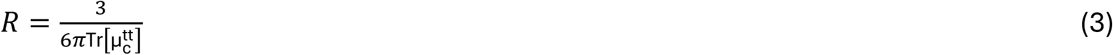

where *µ* ^tt^ stands for translational-translational block of the rigid body grand mobility matrix. We set the size of a bead representing single aminoacid residue to 4 Å, which is optimal based on the previous study (66). We applied GRPY method in its PyGRPY implementation (65, 67, 68).

To obtain a reasonable approximation of α- and β-tubulin hydrodynamic radius, we fed PyGRPY with AlphaFold structures (69, 70) of these proteins: α-tubulin (uniprot: B7Z1K5), and β-tubulin (uniprot: P07437). The obtained hydrodynamic radii are 34.2 Å for α-tubulin, and 28.5 Å for β-tubulin. These values map to friction coefficients in LAMMPS equal to 39.1 and 32.6 g mol^−1^fs^−1^ , respectively.

Interestingly, we found that these friction values are too low to match the slope of MT extreme tip radial distance Δρ observed in MD between 2-4 µs, so we increased them 5-fold. The mismatch is likely due to protein internal friction (71, 72).

### 2.5. Coarse-grained model

We used the all-atom simulations between 2 and 4 µs to parametrize the potentials between CG beads. We adopted a bottom-up approach, specifically Iterative Boltzmann Inversion (IBI) (73–75), in which we iteratively adjusted CG potentials based on comparison of histograms from CG BD simulations (59–61) started at configurations from 2 µs of MD, and ground truth histograms from the all-atom MD simulations. As potentials for the first iteration of IBI, we used Direct Boltzmann Inversion (DBI) potentials.

We introduced potentials governing the following degrees of freedom: (i) longitudinal intra- and (ii) interdimer distances, (iii) αα-, (iv) ββ-, and (v) seam (αβ) lateral distances, (vi) short and (vii) long diagonal distances, (viii) αβα- and (ix) βαβ-longitudinal angles, and (x) longitudinal dihedral angles

Including these interactions in the CG model, we were able to find potentials approximately matching the corresponding histograms, but the global behavior of MTs, specifically the direction of curving, was not matched due to two issues. Firstly, as CG beads in our model present spherical symmetry, in contrast to tubulin monomers which they represent, bending inwards, outwards or sidewards were all enabling the PFs to perform the angular relaxation. However, in MD simulations PFs present exclusively outward bending (Video S1) Secondly, even for PFs bending outwards, we observed substantial indentation of their bottommost segments, unobserved in the MD simulations Thus, we accounted for these discrepancies in our model by two extra terms in the potential energy expression: (xi) dihedral interaction engaging the center of the base of cylinder enclosed by the MT, the lowest bead of a given PF and any pair of consecutive beads along that PF, which leads to an inwards-outwards asymmetry; and (xii) cylindrical repulsive wall inside the MT to prevent the lower parts of PFs from inwards bending. These complete the set of 12 interactions governing our CG model (Fig. S2). The cylindrical wall was centered inside the MT, with a radius of 90 Å, and interacted with tubulin monomers via repulsive harmonic potential of force constant 15 kcal mol^-1^ Å^-2^ and cutoff tubulin-wall distance 10 Å.

For each type of interaction, we assumed that each dimer composing the MT is equivalent, except for angles and dihedral angles, where the histograms were averaged over all triples/quadruples that lie at the tip (4 topmost monomer layers), as we deem them behaving mostly as individual tubulins, i.e., uncoupled from the MT lattice.

We obtain the first approximation of interaction potential V(^1^)(q) as DBI:

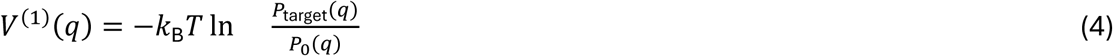

where *q* is a given degree of freedom, *k*_B_ is Boltzmann constant, *T* is temperature, *P*_target_(*q*) is averaged histogram from the reference MD simulations, and *P*_0_(*q*) is the histogram assuming no interactions between the particles, which corrects for the degeneracy associated with a given degree of freedom. For distances:

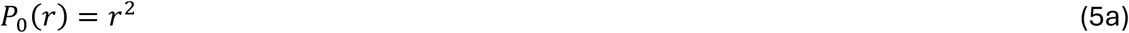

For angles:

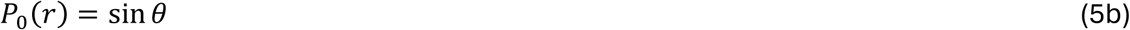

and for dihedral angles:

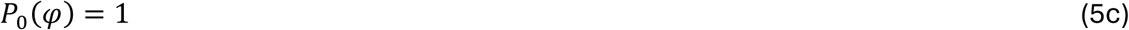

At this point, to avoid any assumptions about the potential shapes, we represented them in a tabular form.

In IBI procedure (73–75), DBI potentials are iteratively improved in the following way:

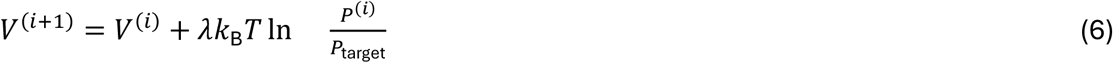

where λ ∈ (0, 1] is a learning rate, V^(i)^ is potential used in i-th round of BD simulations (V(^1^) comes from DBI), which results with P^(i)^ histogram. We shifted the histograms by their minimum nonzero value to avoid computing logarithm of 0. We started IBI from learning rate λ = 0.1 and decreased it to 0.02 and 0.01 along the way. We performed IBI using in-home scripts wrapping LAMMPS (63).

After 60 iterations, we achieved sufficient agreement between corresponding distributions from CG BD and MD (Fig. S3). However, the tabular potentials obtained with IBI were very noisy (Fig. S4).

Furthermore, for lateral and diagonal interactions, the tabular potentials were unstable at large distances, as they were poorly sampled in finite-time MD simulations. For larger numerical stability and to reduce overfitting, we fitted the final IBI tabular potentials with analytic models.

The harmonic potential was used for longitudinal bonds and angles, and for outwards-biasing dihedral angles:

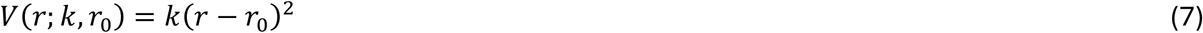

Morse potential was used for lateral and diagonal interactions:

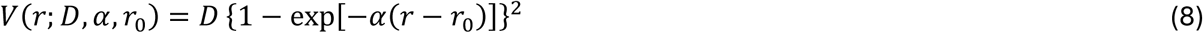

and cosine (dihedral harmonic) was used for longitudinal dihedral angles:

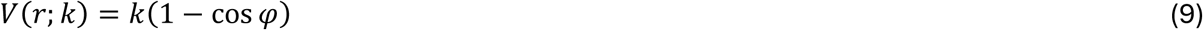

All the final analytic potentials driving our CG model are presented in Fig. S5.

Starting from the final (4 µs) configurations of the MD simulations , five independent 40-µs CG BD simulations for each nucleotide state were conducted. For analysis of the trajectories, we used MDAnalysis (76–78) and in-home scripts.

### 2.6. Cryogenic electron tomography

We used cryo-ET images of MTs initiated from A-sub tubules of alga flagellar axonemes kindly shared by prof. J. Richard McIntosh. The details of sample preparation and tilt series collection were previously reported in the paper by McIntosh et al. (19). We used 3dmod software (79), specifically the slicer window, to trace the MT tips in the tomograms following the protocol described in detail in McIntosh et al.’s paper (19). GTP-MT (polymerizing) samples were obtained at 2 mg/ml tubulin concentration at 37 °C, whereas the GDP-MT (depolymerizing) ones were obtained by subsequent dilution.

From the tomograms, we extracted MT tips’ coordinates. We transformed them to cylindrical coordinates after defining the MT major axis as an average of the vectors pointing from the bottotmmost PF points to the bottommost-but-one ones. The radial vector of each PF was defined as the projection of the vector pointing from the bottommost point to the topmost point to the subspace orthogonal to the MT major axis. The sign was chosen so that the center of the MT lumen would yield negative Δρ. We calculated Δρ and Δz in such defined cylindrical coordinates.

To focus on the extreme MT tips, we included only the segments that deviated by more than 5° from the MT major axis over 10 consecutive line segments. Then, we trimmed the PFs to match the length of those modeled in our simulations to mitigate differences arising from varying PF lengths. The values of PF curvature in a function of distance from the MT tip were adapted from the McIntosh et al.’s paper directly (Fig. 5J and 7H there) (19).

### 2.7. Statistical analysis

We performed statistical analysis of MD trajectories to determine statistically significant differences between bond lengths, angles, and dihedral angles in GDP- and GTP-MT tips. To do that, for each parameter of interest that we deem locally equilibrated (not evolving in time), we computed a time series of its average over all equivalent dimer/trimer/tetramer copies between 2 and 4 µs. As these time series are autocorrelated, we computed their effective sample sizes (80, 81) dividing the total time by the autocorrelation time. The latter we estimated by a time value at which autocorrelation of a given observable drops below 1/e. Using such calculated effective sample sizes, we performed Welch’s t-test (82), which in contrast to the standard t-test does not rely on assumption of homoskedasticity. Results are presented in Table S2.

For observables exhibiting time evolution, we used a different approach, because their behavior exhibits autocorrelation due to the trend that has nothing to do with the sample size. We first performed linear fits to the time series between 2 and 4 µs. Then, we subtracted linear fits from the time series to isolate the noise. Then, we computed the autocorrelation time and effective sample size for the noise time series. Based on these, we computed standard errors of the linear fits which are heteroskedasticity and autocorrelation robust (HAC, Newey-West) (83) using a number of lags equal to the autocorrelation time lag and without small sample correction. The resulting slope coefficients and intercepts were tested for equality (GDP vs. GTP) with Welch’s t-test (82), using sample sizes defined by analysis of error autocorrelation. Results are presented in Table S3.

For angles and dihedrals in distinct PF clusters separately, we additionally computed spatial correlation relaxation timescale of longitudinal angles and dihedrals. Surprisingly, even the nearest lateral neighbors were correlated below 1/e, so we computed effective sample size for each protofilament as described above, through autocorrelation decay for individual protofilament. We performed Welch’s t-tests for equality of means using the resulting effective sample sizes (Tabs. S4-S11).

## 3. RESULTS

### 3.1. Emergent dynamics of GTP- and GDP-MT tips from all-atom MD simulations

To evaluate how the tips of GTP- and GDP-bound MTs relax, we started by analyzing their evolution during all-atom MD simulations (Video S1). The MT tips consisted of 14 PFs each built of 8 α-β-tubulin heterodimers and forming a hollow cylinder with a 3-monomer pitch. During the MD simulations, the MT tips curved outwards, breaking some of the lateral bonds between the PFs (Fig. 1a). This process led to the formation of clusters of PFs. For GDP-MT, 5 clusters with composition of 3-3-3-3-2 were observed, while GTP-MT formed 4 clusters with composition of 4-4-3-3 (Fig. 1b). The seam-composing PFs in both nucleotide types dissociated, indicating weaker heterotypic lateral interactions compared to homotypic ones.

**FIGURE 1.**
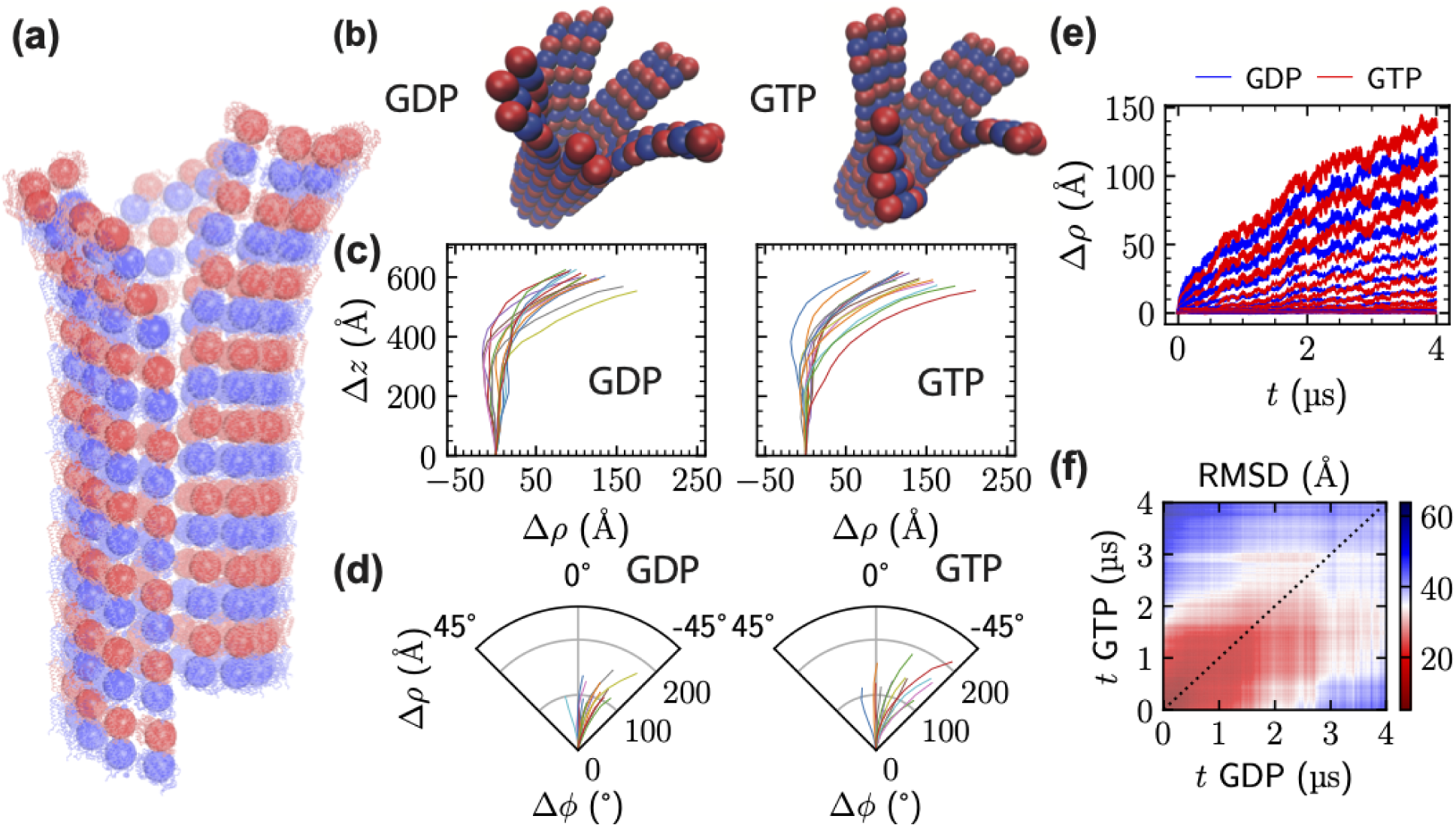
GTP- and GDP-MT plus-end tips are unstable in tubular form and present similar relaxation dynamics in all-atom MD simulations. (a) Overlay of all-atom and CG representations of the MT tip at 0.4 µs of simulation. CG beads representing tubulin monomers (large spheres) are placed at the geometric centers of their C_α_s. Blue beads denote α-tubulin, whereas the red ones denote β- tubulin. (b) CG representation of the GDP- and GTP-MT plus-end tips at 4 µs of MD simulations. (c) Projections of PFs onto their individual radial planes at 4 µs of MD simulation for GDP- and GTP-MT tips. (d) Projections of PFs onto the plane perpendicular to the MT axis at 4 µs of MD simulation for GDP- and GTP-MT tips. (e) Time evolution of radial distance of each tubulin layer from the MT wall Δρ averaged over PFs (layers further from the plus-end MT tip present smaller Δρ values) for GDP- (blue) and GTP-MT (red). Topmost layers 1-3 are plotted with the thickest lines. Layers 4-9 are plotted with intermediate linewidth. Remaining layers are plotted with the thinnest lines. (f) Heat map of pairwise RMSD of GDP- vs. GTP-MT tips computed between the centers of geometry of the C_α_s of corresponding tubulin monomers. Dashed black line denotes the principal diagonal.

Analysis of evolution of the MT tips in cylindrical coordinate system (radial distance Δρ, azimuthal angle Δϕ, and axial distance Δz, Fig. S6; Δ indicates that for each PF, we subtracted the values corresponding to the bottommost, immobile bead to align the PFs), showed that both GDP- and GTP- PFs exhibited similar radial and out-of-plane curving patterns (Fig. 1c-e). All PFs curved exclusively outwards from the MT wall, with indentations in bottom layers below 20 Å (Fig. 1c and Fig. S7). At 4 µs, GTP-PFs exhibited a higher average radial component of curving than the GDP-PFs. GTP-PFs also exhibited greater tangential flexibility (Δϕ = –18.7°, with standard deviation of 16.0°) than GDP-PFs (Δϕ = –15.2°, with standard deviation of 13.2°) at 4 µs (Fig. 1d and Fig. S8).

Layers near the MT tip (layers 1 to 3) continuously curved out with increasing Δρ throughout the simulation (Fig. 1e). Layers further from the tip initially showed minimal Δρ but began increasing after 1.5 µs. Layers close to the MT’s bottom (layers 10 to 16) did not exhibit increasing Δρ, resembling a partially locked zipper. During the first microsecond, GTP-MT tip curved more slowly than GDP-MT one, but GTP-MT tip curved faster after ≈ 0.5 µs (Fig. 1e).

The 4-µs timescale was still insufficient to capture the fully relaxed states of these structures. The root mean squared deviation (RMSD) from the initial tubular structure increased continuously (Fig. S9a). Pairwise RMSD analysis (Fig. 1f) showed that GDP- and GTP-MT tips were comparable during the first 2 µs but diverged thereafter, with GDP-MT tip at 2 µs resembling GTP-MT tip at 1 µs more than GTP- MT tip at 2 µs. To sum up, all-atom MD simulations of MT tips show that both GDP- and GTP-MTs exhibit similar curving behaviors and morphologies, though quantitative differences in radial and tangential PFs curving emerged.

### 3.2. Comparison of inter- and intradimer bond lengths, angles and dihedrals

Although the entire MT tip structure had not equilibrated within 4 µs (RMSD in Fig. S9, Δρ in Fig. 1e), it is possible that distances and angles between tubulin monomers had reached a steady state. To test this hypothesis, we analyzed the evolution of the longitudinal intra- and interdimer distances (Fig. 2a), as well as the longitudinal angles (Fig. 2b and S10), lateral and diagonal distances (Fig. 2c,d), and dihedral angles (Fig. S11).

**FIGURE 2.**
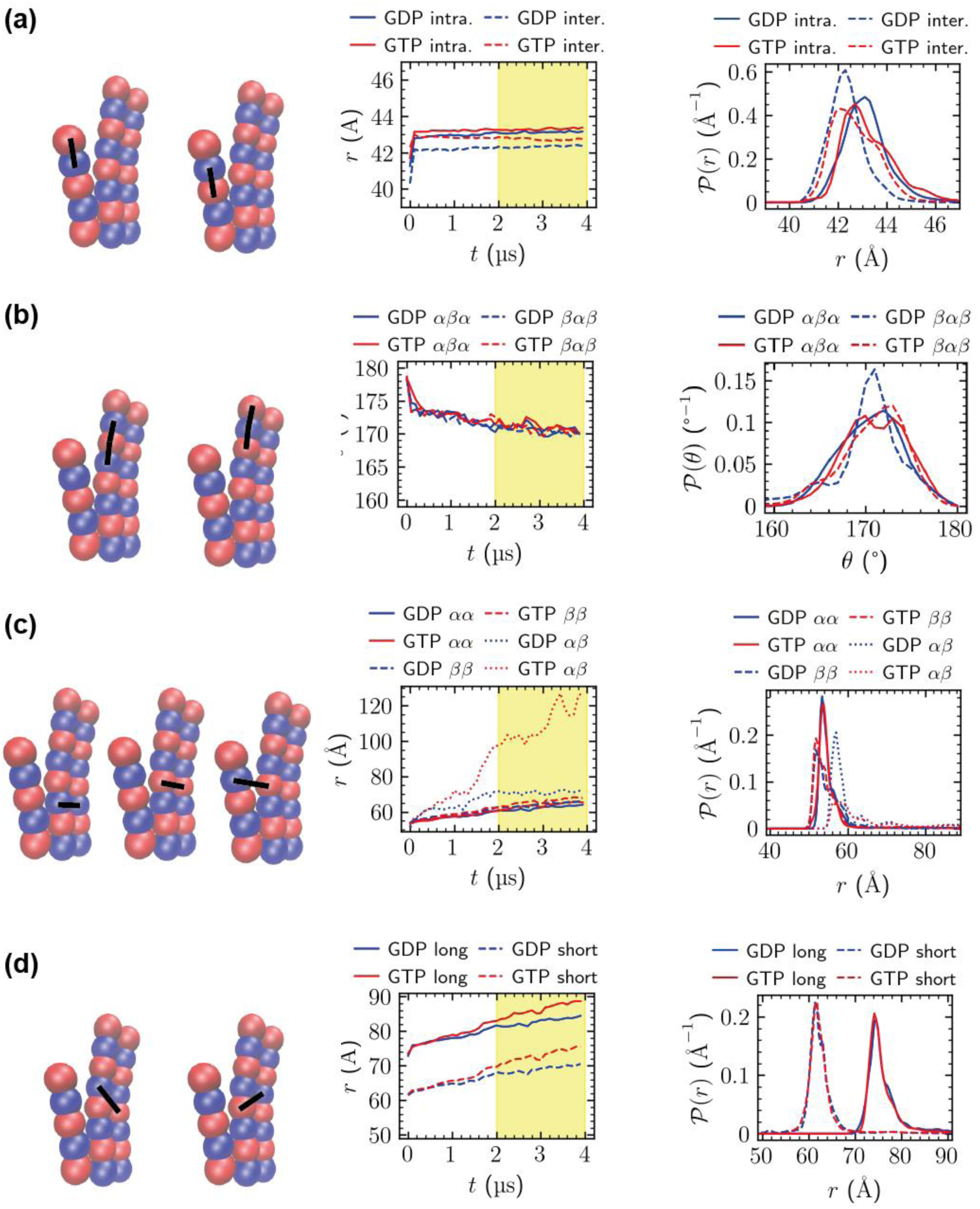
GTP- and GDP-MT plus-end tips present comparable local dynamics in MD simulations. (a) Longitudinal intra- and interdimer bond lengths (example distances marked in the MT tip schemes): time evolution of their values averaged over all symmetry-equivalent pairs throughout the lattice (left), and histograms between 2 and 4 µs (right). (b) Longitudinal αβα and βαβ angles (example angles marked at the MT tip schemes): time evolution of their values averaged over all topmost symmetry-equivalent triples (left), and histograms between 2 and 4 µs (right). (c) Homotypic (αα and ββ) and heterotypic (seam αβ) lateral distances (example distances marked at the MT tip schemes): time evolution of their values averaged over all symmetry-equivalent pairs throughout the lattice (left), and histograms between 2 and 4 µs (right). (d) Short and long diagonal distances (example distances marked at the MT tip schemes): time evolution of their values averaged over all symmetry-equivalent pairs throughout the lattice (left), and histograms between 2 and 4 µs (right). The time traces are plotted with 1000-fold stride to avoid obscuring the trends in the plots, because the noise is much higher otherwise. In all MT tip schemes, blue beads denote α-tubulin, whereas the red ones denote β-tubulin.

Longitudinal inter- and intradimer bond distances were stable throughout the simulation (Fig. 2a and Tab. 1), with intradimer bonds consistently and statistically significantly longer than interdimer bonds. Longitudinal angles between tubulin monomers decreased throughout the simulation, from ≈ 180° to ≈ 170° (Fig. 2b), with minor (less than 0.5°) differences between GDP- and GTP-MT (Tab. 1). However, GDP-MT presented a heavier low-angle tail for the βαβ angle compared to GTP-MT (Fig. 2b), suggesting possible higher radial flexibility, as previously reported (38). Longitudinal dihedral angles near the MT tip predominately acquired values close to 0 and 360° (Fig. S11), indicating a preference for eclipsed (*cis*) configurations. GTP-MT exhibited greater torsional flexibility, with more monomer quadruples between 90 and 270°, compared to GDP-MT, reflecting increased freedom for out-of-plane bending.

**TABLE 1.**
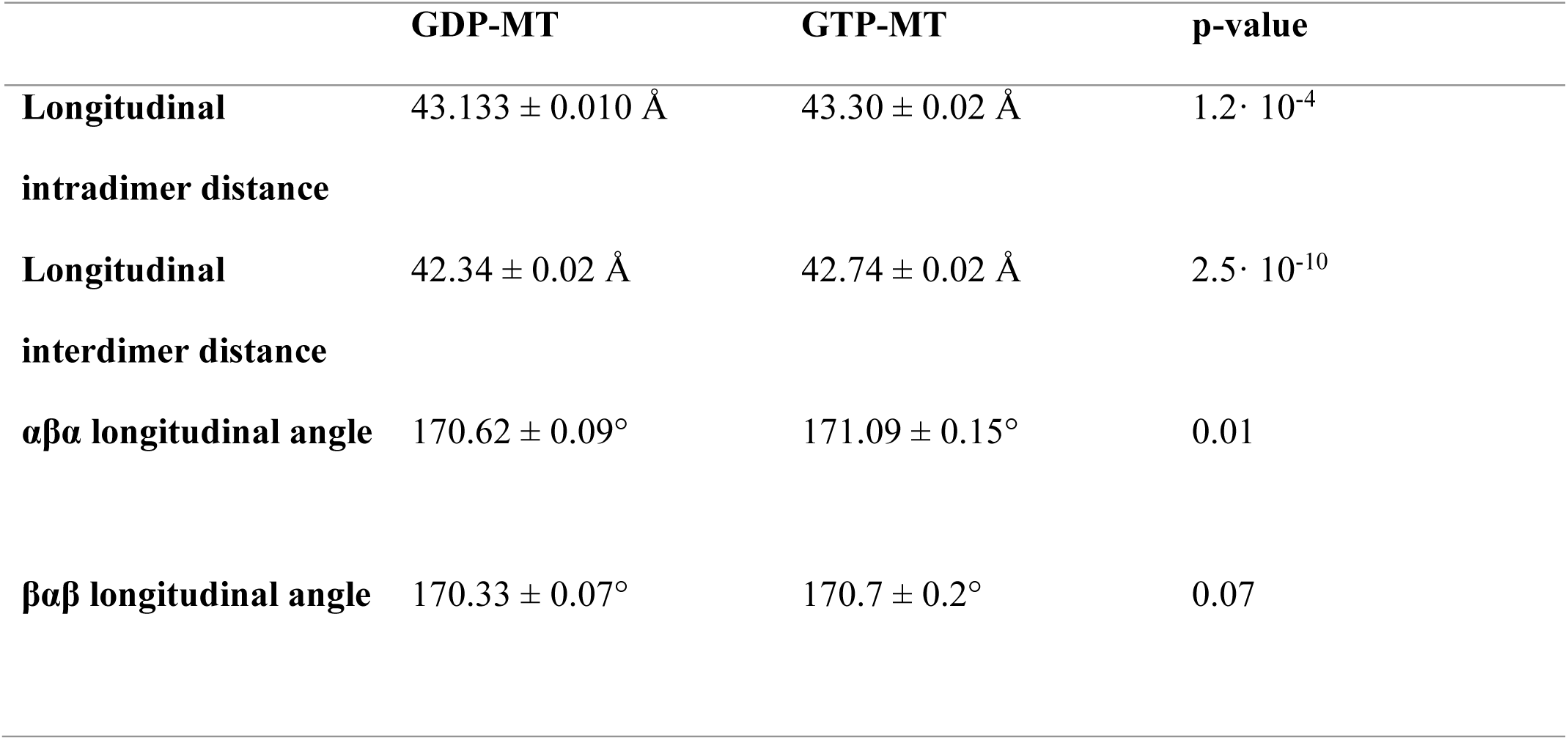
Structural characteristics of GDP- and GTP-MT plus end-tips from all-atom MD simulations between 2 and 4 µs. Longitudinal distances are averaged over the entire MT lattice, while longitudinal angles are averaged over quadruples located at the very tip. The uncertainties associated with the average values denote standard errors of the mean accounting for autocorrelation in the underlying time series. Finally, the p-values are a result of Welch’s t-test used to assess the statistical significance of the observed differences.

Averaging angles and dihedral angles over entire MT tips might obscure the differences in curvature across PF clusters of different sizes. To quantify that effect, we computed histograms of longitudinal angles for PF clusters separately and compared them against each other (Fig. S12). Our analysis shows that there is no clear pattern indicating any coupling between cluster size and curvature. In the GDP-MT case, the differences between various 3-membered clusters are comparable to differences between clusters of sizes 3 and 2. Similarly, in the GTP-MT case, the differences between 3- and 4-membered clusters are comparable to differences between clusters of the same size. We note that it is possible that such coupling emerges for more curved conformations, for which the strain due to bending while keeping neighboring protofilaments close is expected to destabilize lateral clusters. However, we do not observe such coupling in 4 μs of MD.

Lateral distances between tubulin monomers increased steadily throughout the simulation for both GDP- and GTP-MT (Fig. 2c). Histograms of αα-, ββ-, and αβ- (seam) lateral distances showed heavy tails due to monomers liberating and moving away from the MT lattice. Statistical analysis (Tab. 2) revealed a significant difference in the rate of increase for αα-lateral distances between GDP- and GTP-MT, although this was not significant when averaging only over the top layers (p = 0.34, Tab. S3). Also, the ββ-lateral distances presented a significant difference between GDP- and GTP-MT for the whole lattice but not for the top layers (p = 0.22, Tab. S3). This result suggests that significant differences in intermonomer interaction energy between GDP- and GTP-MT concern the vicinity of lateral bond length, likely the dissociation barrier, whereas the long-range interactions are absent. αβ-seam interactions, identified as weak points in the MT structure in previous theoretical (42) and experimental studies (84), showed notable differences between nucleotide states. The time evolution of seam distances differed significantly from homotypic lateral interactions and varied between GDP- and GTP-MT seams.

**TABLE 2.**
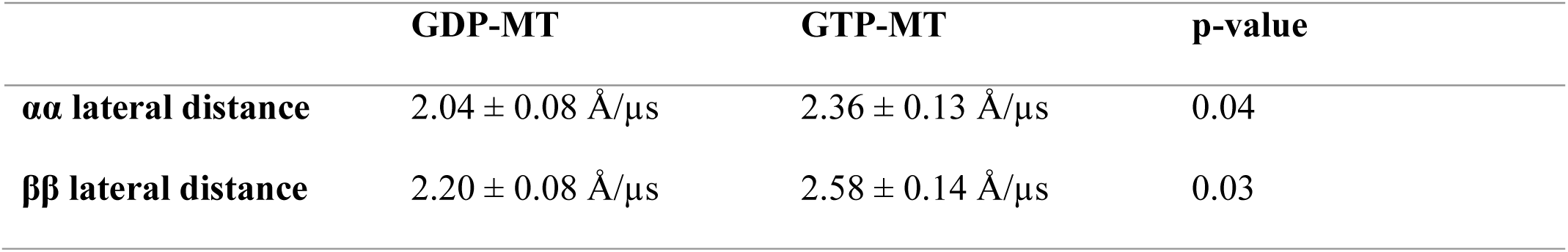
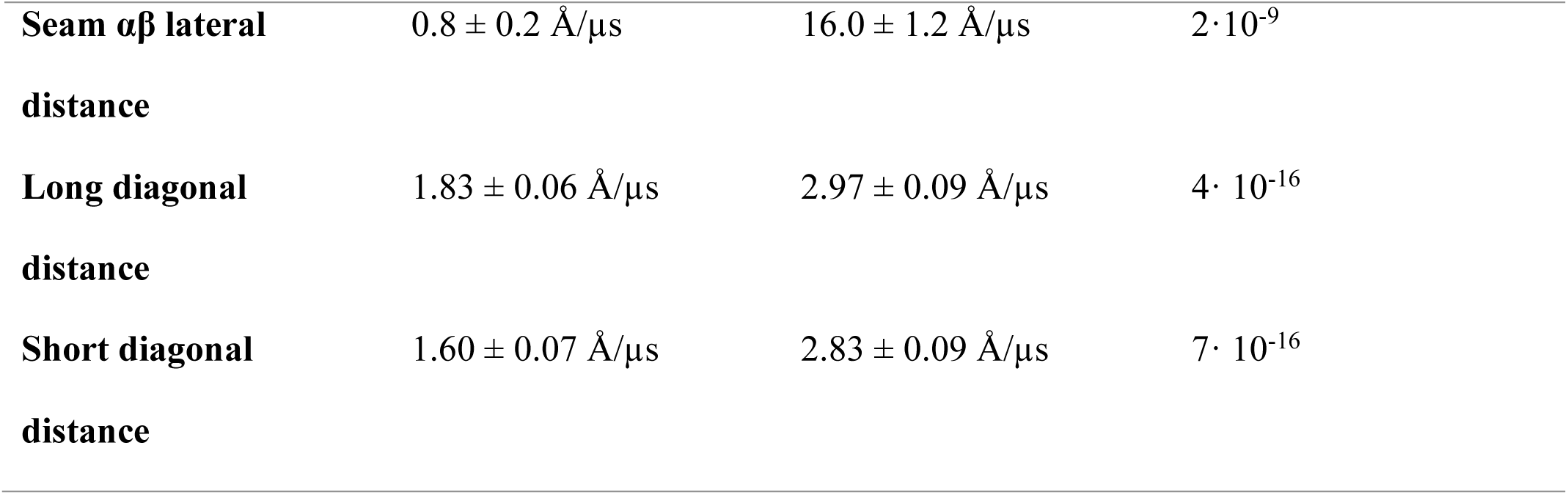
Dynamical characteristics of GDP- and GTP-MT plus end-tips from all-atom MD simulations between 2 and 4 µs. Lateral and diagonal distances are averaged over the entire MT lattice. The uncertainties associated with the rates of change denote standard errors of the linear fits accounting for autocorrelation and heteroskedasticity in the underlying time series. Finally, the p-values are a result of Welch’s t-test used to assess the statistical significance of the observed differences.

GTP-MT had higher average diagonal distances compared to GDP-MT (Fig. 2d): 86 vs. 82.9 Å for long diagonals and 73 vs. 69.1 Å for short diagonals. Statistical analysis (Table 3) of diagonal distances showed significant differences in the rate of change, with faster breakage of diagonal bonds in GTP-MT. However, differences for short diagonals in the top layers were not significant (p = 0.92; Tab. S3).

**TABLE 3.**
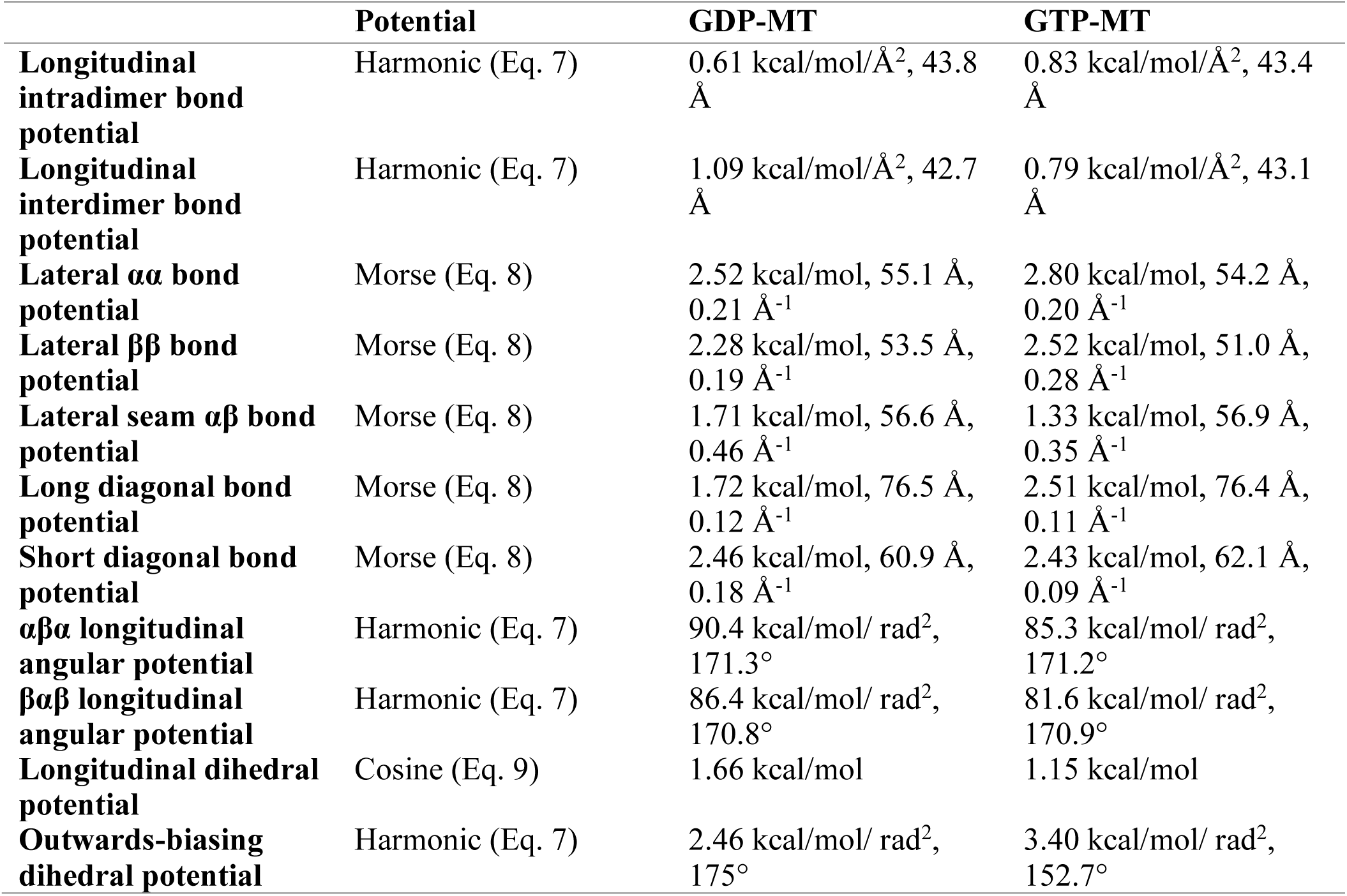
Coarse-grained microtubule model parameters obtained through Iterative Boltzmann Inversion.

|  | Potential | GDP-MT | GTP-MT |
| --- | --- | --- | --- |
| **Longitudinal intradimer bond potential** | Harmonic (Eq. 7) | 0.61 kcal/mol/$\text{\AA}^2$, 43.8 $\text{\AA}$ | 0.83 kcal/mol/$\text{\AA}^2$, 43.4 $\text{\AA}$ |
| **Longitudinal interdimer bond potential** | Harmonic (Eq. 7) | 1.09 kcal/mol/$\text{\AA}^2$, 42.7 $\text{\AA}$ | 0.79 kcal/mol/$\text{\AA}^2$, 43.1 $\text{\AA}$ |
| **Lateral $\alpha\alpha$ bond potential** | Morse (Eq. 8) | 2.52 kcal/mol, 55.1 $\text{\AA}$, 0.21 $\text{\AA}^{-1}$ | 2.80 kcal/mol, 54.2 $\text{\AA}$, 0.20 $\text{\AA}^{-1}$ |
| **Lateral $\beta\beta$ bond potential** | Morse (Eq. 8) | 2.28 kcal/mol, 53.5 $\text{\AA}$, 0.19 $\text{\AA}^{-1}$ | 2.52 kcal/mol, 51.0 $\text{\AA}$, 0.28 $\text{\AA}^{-1}$ |
| **Lateral seam $\alpha\beta$ bond potential** | Morse (Eq. 8) | 1.71 kcal/mol, 56.6 $\text{\AA}$, 0.46 $\text{\AA}^{-1}$ | 1.33 kcal/mol, 56.9 $\text{\AA}$, 0.35 $\text{\AA}^{-1}$ |
| **Long diagonal bond potential** | Morse (Eq. 8) | 1.72 kcal/mol, 76.5 $\text{\AA}$, 0.12 $\text{\AA}^{-1}$ | 2.51 kcal/mol, 76.4 $\text{\AA}$, 0.11 $\text{\AA}^{-1}$ |
| **Short diagonal bond potential** | Morse (Eq. 8) | 2.46 kcal/mol, 60.9 $\text{\AA}$, 0.18 $\text{\AA}^{-1}$ | 2.43 kcal/mol, 62.1 $\text{\AA}$, 0.09 $\text{\AA}^{-1}$ |
| **$\alpha\beta\alpha$ longitudinal angular potential** | Harmonic (Eq. 7) | 90.4 kcal/mol/ $\text{rad}^2$, 171.3$^\circ$ | 85.3 kcal/mol/ $\text{rad}^2$, 171.2$^\circ$ |
| **$\beta\alpha\beta$ longitudinal angular potential** | Harmonic (Eq. 7) | 86.4 kcal/mol/ $\text{rad}^2$, 170.8$^\circ$ | 81.6 kcal/mol/ $\text{rad}^2$, 170.9$^\circ$ |
| **Longitudinal dihedral potential** | Cosine (Eq. 9) | 1.66 kcal/mol | 1.15 kcal/mol |
| **Outwards-biasing dihedral potential** | Harmonic (Eq. 7) | 2.46 kcal/mol/ $\text{rad}^2$, 175$^\circ$ | 3.40 kcal/mol/ $\text{rad}^2$, 152.7$^\circ$ |

Collectively, these results show that inter- and intradimer longitudinal bond lengths, homotypic lateral bond lengths, and longitudinal angles are comparable between the nucleotide states. In contrast, larger differences were observed in the interactions along the diagonals of parallelograms composed of tubulin quadruples tiling the MT wall, in the interactions at the seam, and in the tangential flexibility of PFs.

These findings suggest that the GTP-MT lattice exhibits high dynamism, which aligns with the understanding that GTP-MT tips are also curved (19, 31–34, 42, 43), both radially and tangentially (out-of-plane).

### 3.3. Extended dynamics and equilibration of GTP- and GDP-MT tips using CG-ing

To evaluate the structural changes of GDP- and GTP-MT plus-end tips beyond 4 µs all-atom MD simulations, a CG model was parameterized and utilized (Tab. 3). The PFs’ clustering observed during the MD simulations persisted in the CG BD simulations: GDP-MT tip retained 5 clusters with a 3-3-3-3- 2 composition, while GTP-MT tip retained 4 clusters with a 4-4-3-3 composition (Fig. 3a), both presenting curvatures increasing over time. To determine whether different cluster numbers stem from underlying differences in interactions or result simply from underlying stochasticity of the MT relaxation, we performed 20 0.2-μs-long replicas of CG BD simulations starting from tubular GDP- and GTP-MT. Average cluster numbers in GDP- (4.75 ± 0.14) and GTP-MT (5.00 ± 0.13) did not differ statistically significantly (p=0.18, Fig. S13), confirming remarkable similarity between GDP- and GTP- MT plus-end tip relaxation.

**FIGURE 3.**
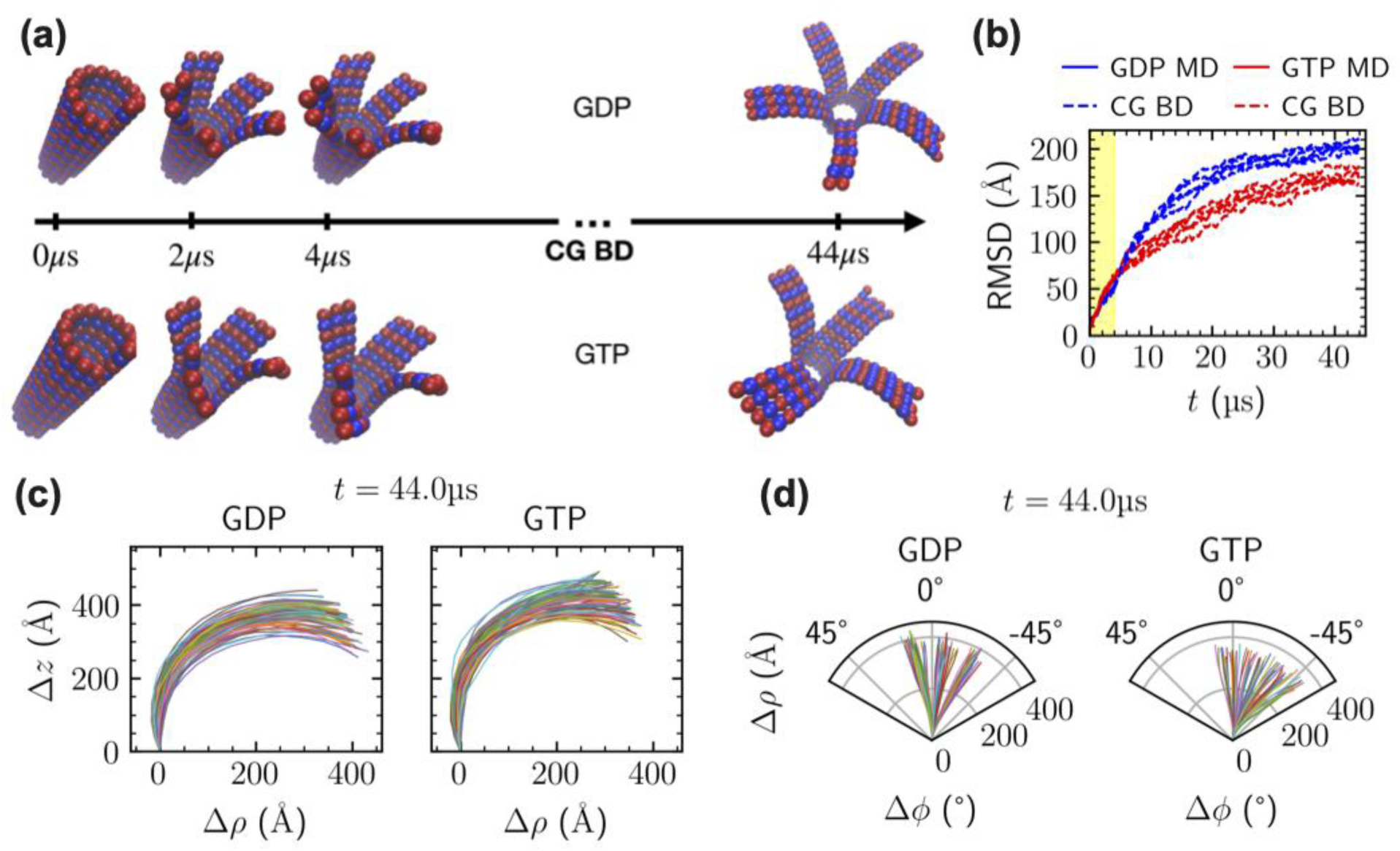
Time evolution of MT plus-end tip structures for tens of microseconds. (a) CG representation of GDP- and GTP-MT tip structures at the onset of MD, at 2 and 4 µs of MD, and at 44 µs of CG BD. Blue beads denote α-tubulin, whereas the red ones denote β-tubulin. (b) Plot of RMSD vs. time for 5 CG BD replicas, computed relative to the initial, cylindrical configuration. (c) Projections of PFs onto their individual radial planes at 44 µs of CG BD simulation. (d) Projections of PFs onto plane perpendicular to the MT axis at 44 µs of CG BD simulation.

Unlike the MD simulations, the CG BD simulations showed signs of equilibration, as indicated by the saturation of RMSD relative to the initial tubular conformations at about 40 µs (Fig. 3b) and by pairwise RMSD demonstrating that the configurations sampled during the last 10 µs of the CG BD simulations were similar between nucleotide states (Fig. S14).

At the end of the CG BD simulations, both GDP- and GTP-MT tips exhibited increased bending compared to the end of the all-atom MD simulations (Fig. 3c vs. Fig. 1c). Specifically, the average radial distance of the MT tip from the MT wall increased from Δρ = 117.3 Å to 379.6 Å for GDP-MT and from 135.6 Å to 332.6 Å for GTP-MT. This suggests a substantial increase in radial bending, with GDP-MT tip presenting higher radial coordinate. The PFs in the CG BD simulations continued to flare, with lower layers also curving outward. Additionally, GTP-PFs displayed more tangential dispersion compared to GDP-PFs (Fig. 3d).

The radial distance ρ averaged over MT tips during the CG BD simulations continued to evolve smoothly beyond the all-atom MD (Fig. S15). Small differences in long-timescale relaxation between GDP- and GTP-MT tips were influenced by the nucleotide state rather than initial cluster composition (Fig. S15). Collectively, results from analysis of the CG BD simulations show that GDP- and GTP-MT tips exhibit high structural resemblance within timescale beyond the one achievable by all-atom MD simulations.

### 3.4. Comparison of simulated GTP- and GDP-MT tips with experimental images

To evaluate how the CG BD relaxed MT plus-end tip structures compare with MT tip morphologies detected experimentally, the modeling results were validated through comparison with cryo-ET structures of polymerizing and depolymerizing MT tips (19) (Fig. 4a). Most of the cryo-ET MT tips displayed curvature consistent with the mean curvature from our CG BD simulations (Fig. 4b).

**FIGURE 4.**
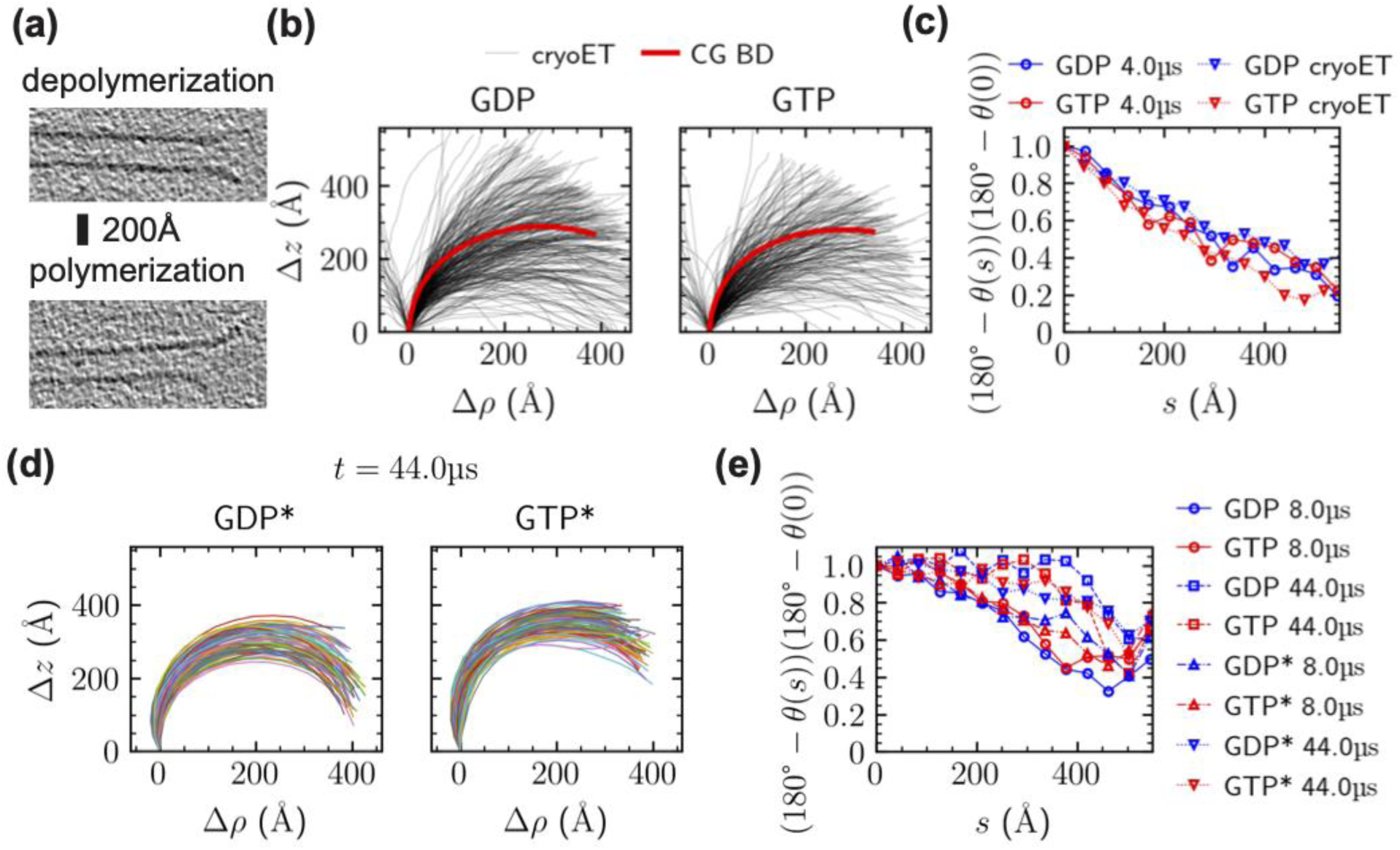
Validation of MD and CG BD simulations against cryo-ET MT plus-end tip structures. (a) Example slices through the end of polymerizing and depolymerizing MT tips. (b) Projections of cryo-ET MT tips onto their individual radial planes. PFs are trimmed from the bottom based on deviation from straight conformation, and from the top by setting their length equal to the length of MT tips from simulations. (c) Dependence of relative curvature of the PFs on the distance from the tip *s* from MD simulations and cryo-ET. (d) Projections of PFs onto their individual radial planes at 44 µs of CG BD simulations with top-down corrected angular potentials. (e) Relative curvature of a PF as a function of distance from the tip *s* at different times of CG BD simulations with (asterisks) and without top-down corrected angular potentials.

Simulated and experimental GDP- versus GTP-MT tips did not differ qualitatively in their curvature.

Cryo-ET data exhibited the linear dependence of PF curvature on the distance from the MT tip (Fig. 4c and Fig. S16a), a pattern not captured by existing MT models (19). Our MD simulations exhibited the same linear relationship. Although the absolute curvature values in the experimental tips are higher (Fig. S16a), when normalized relative to the MT tip, the curvature profiles are comparable (Fig. 4c).

To address the potential underestimation of equilibrium curvature in our CG model, the angular potentials were adjusted by shifting their minima by 3° (GDP* and GTP* models), resulting in more curved PFs (Fig. 4d). In the CG BD simulations, a similar linear relaxation pattern emerged until the tip reached equilibrium curvature (Fig. 4e and Fig. S16b). Once equilibrium at the tip was established, the neighboring layers of the tip adopted a curvature that minimized their angular potential energy, causing the curvature-distance relationship to level off.

In summary, data from our model align well with experimental cryo-ET images, validating its accuracy in capturing the curvature characteristics and relaxation patterns of MT tips in the two nucleotide states. This agreement demonstrates the robustness of our CG BD simulations in capturing the dynamics of GTP- and GDP-MT tips and supports the idea that GTP- and GDP-MTs exhibit comparable in-plane and out-of-plane bending, further highlighting a linear dependence of curvature on distance from the tip.

## 4. DISCUSSION

### 4.1. GTP- and GDP-MT tips do not differ substantially in curvature and lateral interactions

Our study addresses the ongoing debate about the mechanisms underlying the structural dynamics of the MT plus-end tip, particularly within the context of the allosteric and lattice models (32). Both models attribute MT instability to tip curvature relaxation due to decreased lateral interactions, with or without tubulin conformation changes. The allosteric model suggests that GDP-tubulin, being more bent than GTP-tubulin, causes strain in the MT lattice, promoting outward bending of PFs (22–24). In contrast, the lattice model links increased bending to weaker lateral interactions within the MT lattice (25, 26, 38).

Our findings challenge these models by revealing similar PF flaring, lateral interactions, and intrinsic curvatures between GDP- and GTP-bound MT tips (Figs. 2–4). Previous work has already questioned the validity of the allosteric and lattice models, suggesting that a combination of both mechanisms might explain MT instability and outward PF bending (85). Additionally, GDP- and GTP-bound MT tips are reported to exhibit similar intrinsic curvatures (19, 27–33, 35–38), which aligns with our findings. The assumption of distinct curvatures between GDP- and GTP-MTs, which was central to several previous theoretical models (86–88), is contradicted by our all-atom MD data. Our results challenge the allosteric model’s assumption that differences in intrinsic curvature between GDP- and GTP-tubulins drive bending (Fig. 2b and Table S2), and is consistent with recent studies (36, 38). Regarding the lattice model, our simulations show that homotypic lateral interactions are also comparable between GDP- and GTP-MTs (Fig. 2c). Interestingly, GTP-MTs exhibit slightly stronger lateral bonds than GDP-MTs, with αα-lateral interactions slightly stronger than ββ-lateral interactions, in line with the stabilization of the M-loop in GTP-bound α-tubulin (38, 89). This suggests that lateral interactions alone cannot explain the primary determinant of MT tip stability.

The seam, a weak interface of the MT cylindrical structure, has been shown to break during simulations (42, 84). In our simulations, the seam breaks in both GDP- and GTP-MT tips (Figs. 1b, 2c), with dissociation energies ∼ 29% weaker in GDP-MTs and ∼50% weaker in GTP-MTs compared to homotypic dissociation energies. Differences between GTP- and GDP-lattices primarily arise from interactions along the MT lattice diagonals, with dissociation energies of 1.72 kcal/mol for long- diagonal interactions in GDP-MTs and 2.51 kcal/mol in GTP-MTs, and the coupling between longitudinal tubulin dimers.

Our study focuses on the extreme plus-end tip, and it is plausible that changes in tubulin structural dynamics between GTP- and GDP-MTs may be more pronounced closer to the bulk. The GTP cap can extend over many heterodimers, influencing overall stability (90), and there may be a transition point between the flaring plus-end tip and the bulk MT lattice where GTP/GDP composition significantly affects MT dynamics. Interestingly, no dissociation of longitudinal interdimer bonds was observed in our MD simulations (Fig. 2a). Future work could explore how GTP hydrolysis affects the stability of longitudinal interfaces. Additionally, the angular potentials between tubulin triples in our model were harmonic, neglecting potential effects of low-angle tails in the angle histograms (Fig. 2b). Investigating the statistical significance of these tails and their potential effects on long-term MT tip dynamics will be an important direction for future studies.

### 4.2. Linear curvature-tip distance relation in MT tip relaxation

A linear relation between the PF curvature and distance from the tip has been reported in cryo-ET study of both polymerizing and depolymerizing MT tips (19). Linear fits to the MD and cryo-ET datapoints (Fig. 4c) yield the slope of 0.11 (0.12) and 0.08 (0.06) %/Å, for GDP (GTP), respectively. This observation raises the question of whether the linearly varying curvature is an equilibrium property or rather a transient feature of MT relaxation. While the presence of this relationship in cryo-ET data suggests it may be an equilibrium property, it is difficult to understand why tubulin dimers close to the tip would not relax to their equilibrium curvature in experiments but instead remain stable at some intermediate value. As such, although the linear relationship between curvature and distance from the MT tip is observed in both models and experiments, it seems more likely to represent a transient feature during the relaxation process rather than a stable equilibrium property. It is also possible that the faster relaxation in simulations, compared to experiments, lead to the MT tip reaching equilibrium more quickly than observed experimentally. This interpretation is further supported by the curvature-distance relations observed in CG BD simulations, particularly in the top-down corrected models (GDP*, GTP*, see Fig. 4e).

### 4.3. PFs at MT tip form stable clusters

Like in previous all-atom MD simulations of MT tips dynamics (42), during the initial stages of our all-atom MD simulations, the PFs remained laterally bound into a few clusters (5 for GDP-MT and 4 for GTP-MT tip, see Fig. 1b). The number and size of clusters are known to be stochastic (42), but the GTP-MT exhibit clusters of larger size (34), which agrees with our results (Fig. 1b and Fig. S13). Our CG model predicts that the clusters remain stable throughout the 40-µs CG BD simulations. However, the number and composition of clusters do not determine the final MT tip state observed at the end of CG BD simulations, which proves that the dynamics of flaring is independent of the initial lattice geometry in terms of clusters number and composition, but depends solely on the interaction potentials between tubulins (Fig. S15).

Curving outwards of PF clusters was captured in MD studies (42), and observed experimentally (34, 43). It was recently suggested that the PF clusters are larger and more stable in GTP-MT, which is consistent with our results (34). Moreover, outward-curved sheet-like structures at the growing MT tips were observed in cryogenic electron microscopy (cryo-EM) images (44, 91) and their stability was demonstrated in CG simulations (86). PFs under high magnesium concentration were also shown to form tubes of diameter considerably larger than in MTs (38).

Our study suggests that GDP- and GTP-MT tips exhibit highly similar PF clustering, with only minor deviations such as heavy tails in GDP-MT βαβ-angle distribution (Fig. 2b), higher torsional flexibility of GTP-MT (Fig. S11), and higher probability of large clusters in GTP-MT (Fig. S13). It is possible that such small deviations can still have functional significance. The βαβ-angle heavy tail, originating almost exclusively from the GHI cluster (Fig. S12), may indicate a specific overbending mode with functional relevance. Increased torsional flexibility of GTP-PFs might lead to smaller clusters merging into larger sheets that can stabilize the overall structure.

Our bottom-up model offers a rigorous framework to explore, in the future, the effect of systematic perturbation of lateral interactions away from the all-atom potentials, combined with variations in bending and twisting flexibility, on the transitions between flared and cylindrical tip morphologies, as well as their relationship with other morphologies such as sheets and tubes.

### 4.4. PFs curving: radial and tangential components

The simulated GDP- and GTP-MT tips displayed radial (Fig. 1c and Fig. 3c) and tangential, out-of-plane bending (Fig. 1d and Fig. 3d), in line with experimental data indicating that bending is not unique to GDP-bound MT tips (19, 27–38).

In 4 µs of MD, the GTP-MT tip curved to a somewhat larger extent than its hydrolyzed counterpart, but this trend reversed during the 40-µs CG BD simulations. We also note that in the first 0.8 µs of MD GDP-MT tip also curved slightly faster. These fluctuations and the fact that at the very end of our CG BD simulations the difference in the extent of curving was as small as diameter of a single tubulin monomer, suggest that GDP- and GTP-MT tip structures and dynamics are very similar.

Different from prior CG models of MT tip dynamics (19, 31, 92, 93), our model did not impose restraints on the tangential motion of the PFs. Indeed, the PFs did not curve exclusively in their respective radial planes (Fig. 1d), but part of their motion was out-of-plane tangential swing clockwise looking from above the MT tip. The orientation is the same as the orientation of the helical MT pattern, i.e., starting from the bottommost bead, the *z* coordinate (pointing to the plus-end) increases clockwise (in other words, the helix is left-handed), as in recent simulations (42) and experiments (34).

In all-atom MD simulations, this tangential swing tendency was slightly stronger for GTP-MT tip, agreeing with previous MD simulations (42) but in contrast to a cryo-EM study suggesting the opposite (38). Our CG model reflects this tendency through parametrization of dihedral potentials. Firstly, the dihedral cosine potential has lower maximum at 180° for GTP-MT (Fig. S5e), allowing less correlated arrangement of consecutive longitudinal monomer triples. Secondly, the outward-biasing dihedral potential for GTP-MT has a minimum at 152.7°, as compared to 175° for GDP-MT, causing larger deviation from in-plane curving (Fig. S5f). Yet, the arrangement of PF traces in axial cuts from long CG BD simulations is dominated by the signatures of PF clustering, rather than by the tangential swing (Fig. 3d).

The outward-bias potential responsible for steering the plane of PF curving is approximate and is expected to break down for longer PFs. This is because such defined dihedral angle for outwards bending is ≈ 180° up to the point when the PF bends into a half-circle. When the ram’s horn exceeds the length needed to form a half-circle, the bias starts favoring its inflection to continue the trend of beads closer to tip being more distant from the MT wall. We note that ram’s horns so long to exceed a half-circle length are rarely observed experimentally (19, 31, 33), and not observed in our simulations (Fig. 3c).

To overcome this limitation in future studies, we propose a modification of the model in which the outwards bias potential can be expressed in such a way that it ensures certain handedness of the ram’s horn spiral. For each consecutive triple of beads *(i, j, k)* along the PF, a unit vector *n_ijk_*can be defined that is parallel to a cross product of vectors joining *i-j* and *j-k*. In case of ideally radial ram’s horn, *n_ijk_* is tangential to MT wall and points in the counterclockwise direction. Allowing for some clockwise tangential swing, the vector tilts in direction opposite to the MT major axis; conversely counterclockwise tangential swing tilts *n_ijk_* along the MT major axis. Potential based on histograms of these projections (Fig. S17) might be considered as a generalization of our outwards bias to MTs of arbitrary length.

### 4.5. Construction of the CG model

We used BI method (73–75) to parameterize potentials governing our CG model. In the IBI procedure, the potentials between tubulin monomers were iteratively updated to match the ground truth histograms from all-atom MD simulations between 2 and 4 µs. Friction coefficients for the BD simulations were obtained using hydrodynamic calculations (65), and then increased five-fold to match the slope of Δρ between 2 and 4 µs (Fig. S18), as several factors contributed to the need for adjustment: hydrodynamic interactions necessary for accurately matching macromolecular mobility were neglected (64), the TIP3P water model used in MD simulations underestimates water viscosity (94); the MT structure altered water relaxation and reorientation, thereby affecting dynamic viscosity (72), and intra-protein degrees of freedom were neglected in the CG model, impacting internal friction (72). Interestingly, the internal friction of myoglobin obtained by measuring the rate constant of its conformational change as a function of solvent viscosity was ≈ 4 times greater than the viscosity of water, which accurately matches our 5- fold friction upscaling (71). Our combined parametrization of friction and potentials lead to reasonably smooth behavior of ρ at 4 µs, where the all-atom MD and CG BD curves meet (Fig. S15a).

We based our CG model on the longest continuous trajectory of the MT tip, as its relaxation is a slow, far-from-equilibrium process. Equilibrium properties—such as bond lengths, curvatures, and force constants—are more likely to be reliably captured in a single long trajectory than in several shorter ones, where a greater proportion of time is spent on system relaxation. Our approach is thus complementary to one employed by Igaev et al. (42) , who used multiple 1 μs simulations, and Kalutskii et al., (34), who used multiple 2.5 μs simulations, favoring broader sampling through shorter trajectories.

In contrast to a recent bottom-up study of MT tip dynamics (34), here we did not treat any of the interactions for the CG model as free parameters but directly extracted them from the all-atom simulations. The model provides a general framework for simulations of short MT tips, reducing the description of a complex all-atom MT to a simplified model with only 12 interaction potentials. The selection of the 12 specific interactions is sufficient to generate realistic dynamics of MT tips, including outward curving, tangential swinging, and PF clustering. The parameters of this model were derived from all-atom MD simulations and IBI, though they could still be refined with top-down corrections in future studies.

## 5. CONCLUSION

In this paper, we used a multiscale approach combining all-atom MD and monomer-resolution CG BD simulations to explore the relaxation of cylindrical MT plus-end tips in different nucleotide states. The tubular MT tips were unstable in both nucleotide forms and their relaxation dynamics was comparable, presenting PFs clustering, outward bending and out-of-plane PFs’ curving. Additionally, the PFs exhibited linear dependence of curvature angle on distance from the tip, as experimentally observed (19). The difference between the GDP- and GTP-MT tip relaxation is small and only quantitative, not qualitative. This observation aligns with the cryo-ET structures of polymerizing and depolymerizing MTs, both exhibiting bent, ram’s horn-like tip morphologies (19). These findings support the idea that PF bending is not driving the switch to MT depolymerization, challenging two prominent explanations for MT stability-hydrolysis coupling: allosteric and lattice models. MT hydrolysis-stability coupling likely involves factors beyond the scope of our model, such as conformational changes during GTP hydrolysis, slow conformational changes of tubulins not sampled by our all-atom MD simulation or differences in longitudinal dissociation energies, in particular their coupling to PF curvature.

Overall, our multiscale modeling approach provides insights into MT tip dynamics and supports the notion that MT tip relaxation is not coupled to depolymerization. Importantly, these results apply to the very top of the MT plus-end tip and other structural changes and dynamical features may be pronounced in between these “tipmost” heterodimers and the bulk MT lattice. Future research should focus on refining models to account for these observations and exploring additional factors that may influence MT stability and dynamics, both in the bulk MT lattice and in the region marking the change between the bulk and plus-end tip. The interplay between radial and out-of-plane relaxation modes and the linear dependence of PF bending angle on distance from the tip should also be accounted for in future frameworks of MT dynamics. Our model parameters could be further refined through extending MD simulations or incorporating top-down corrections based on experimental data, but our selection of tubulin interactions is sufficient for capturing PF morphological evolution across timescale of several tens of microseconds. In the future, it would be interesting to apply the model to explore other important aspects of MT relaxation such as the effect of unequal PF length on their clustering (95) or the effect of macromolecular crowding on PF bending (96).

## AUTHOR CONTRIBUTIONS

T.C.B., T.S., G.A.V., and D.B. designed the research; T.S., D.B., J.W., and W. X. performed the research; T.S. analyzed the data; T.S. wrote the first draft; all authors edited the final manuscript.

## ACKNOWLEDGEMENTS

This research was supported by the Department of Energy, Office of Science, Basic Energy Sciences, under Award DE-SC0023318 and by the National Institute of General Medical Sciences of the NIH through Fellowship F32 GM140646 (D.B.). Computational resources were provided by the University of Chicago Research Computing Center, the XSEDE computing cluster and the Center for High Performance Computing at the University of Utah. We thank Prof. J. Richard McIntosh for sharing the cryo-ET data, fruitful discussions and inspiring suggestions.

