## Supplementary Materials for "On the Curvature and Relaxation of Microtubule Plus-end Tips"

**Supplementary Figures**

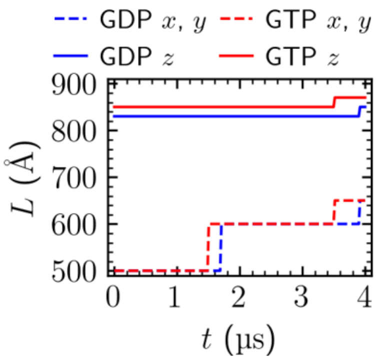


**FIGURE S1.** Time evolution of the simulation box size upon resizing applied in all-atom molecular dynamics simulations of GDP- and GTP-microtubule plus-end tips..

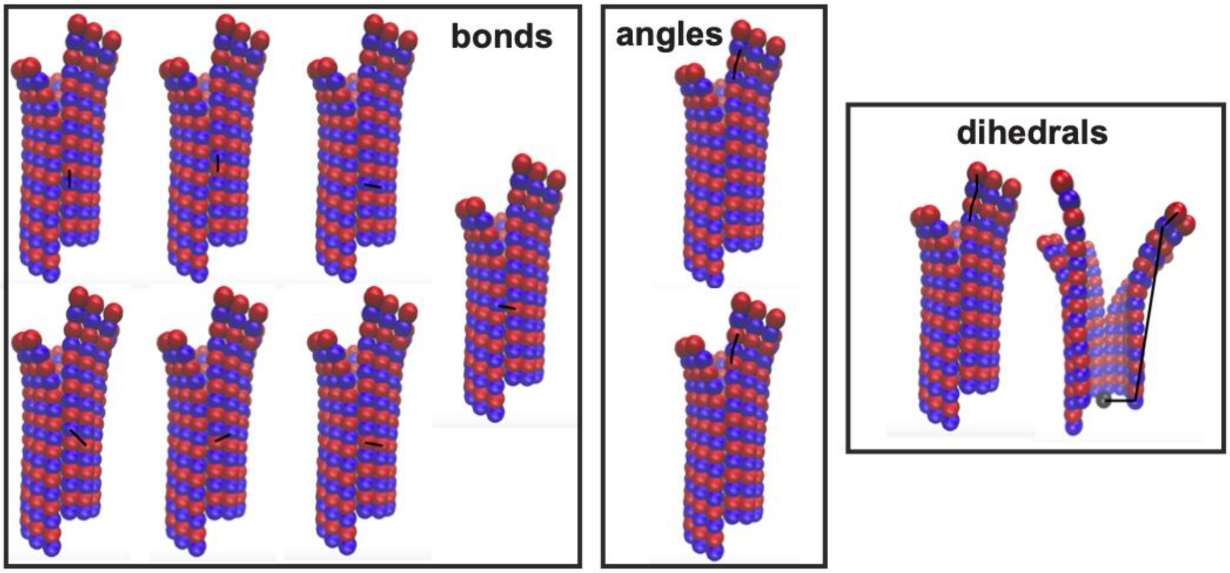


**FIGURE S2.** Monomer-resolution MT plus-end tip CG model is built of 11 interactions: 7 bonds, 2 angles, and 2 dihedral angles. Moreover, inside MT, a repulsive cylinder is placed, that prevents PFs from bending inwards.

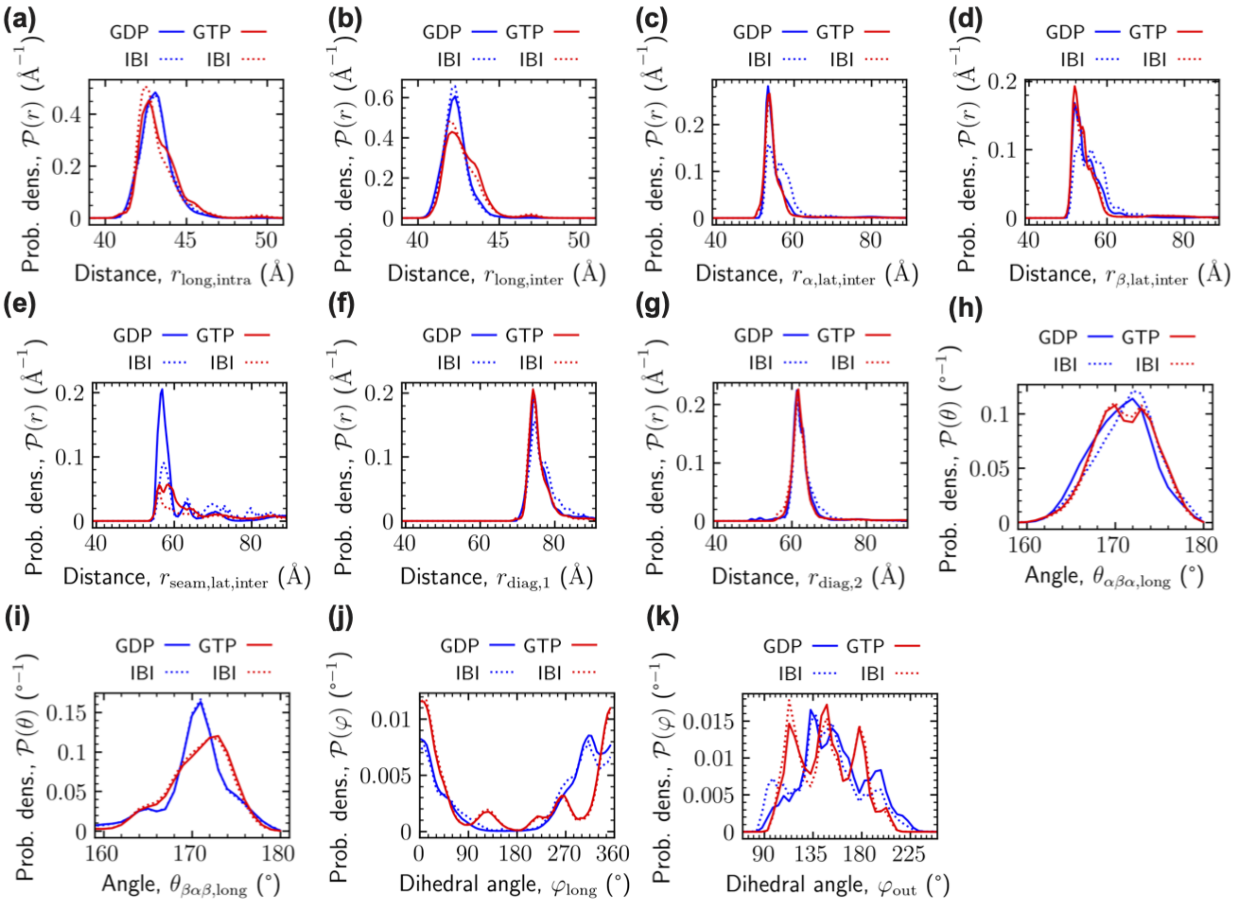


**FIGURE S3.** Comparison of histograms between all-atom MD simulations and final iteration of IBI: longitudinal (a) intra- and (b) interdimer distances; lateral (c) αα, (d) ββ, (e) and αβ (seam) lateral distances; (f) long and (g) short diagonal distances; (h) αβα and (i) βαβ longitudinal angles; (j) longitudinal dihedral angles, and (k) outward bias dihedral angles.

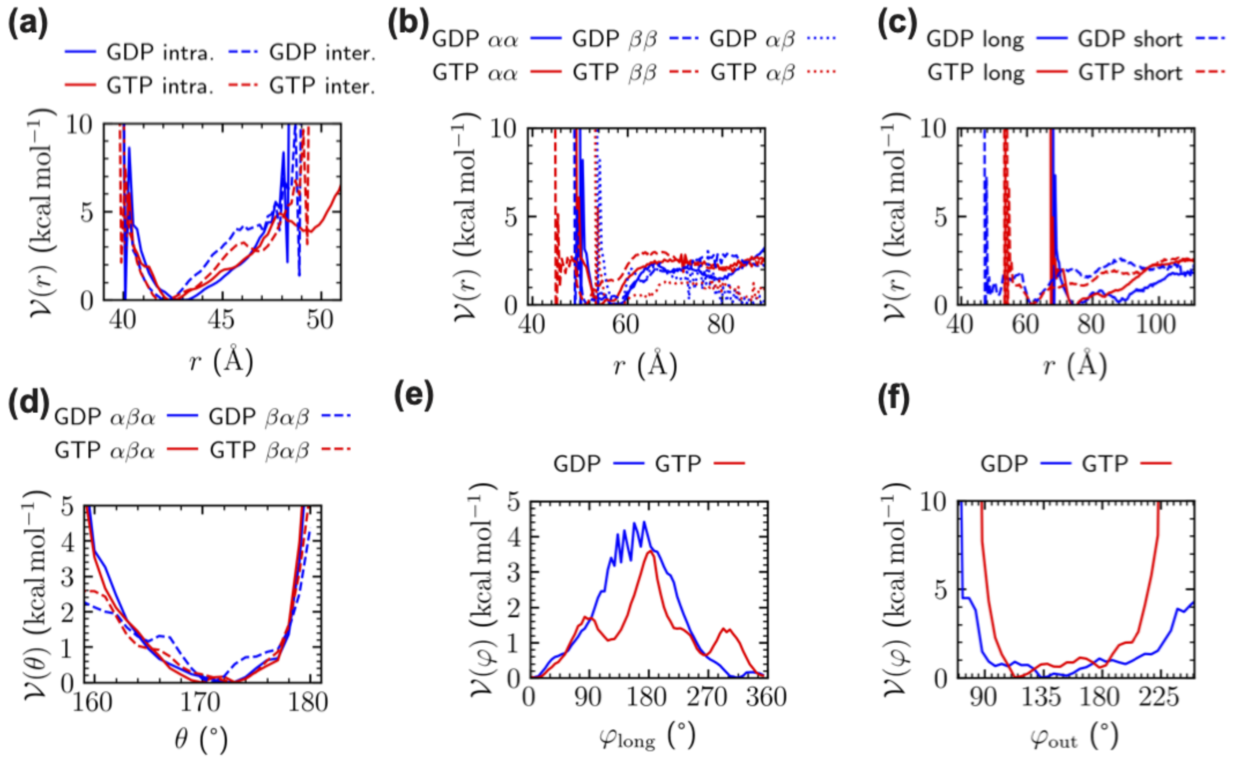


**FIGURE S4.** Comparison of tabular potentials obtained from applying IBI to GDP- and GTP-MT. The analytic potentials from Figure S11 are fits to these tabular potentials. (a) intra- and interdimer distances; lateral (b) αα, ββ, and αβ (seam) lateral distances; (c) long and short diagonal distances; (d) αβα and βαβ longitudinal angles; (e) longitudinal dihedral angles, and (f) outward bias dihedral angles.

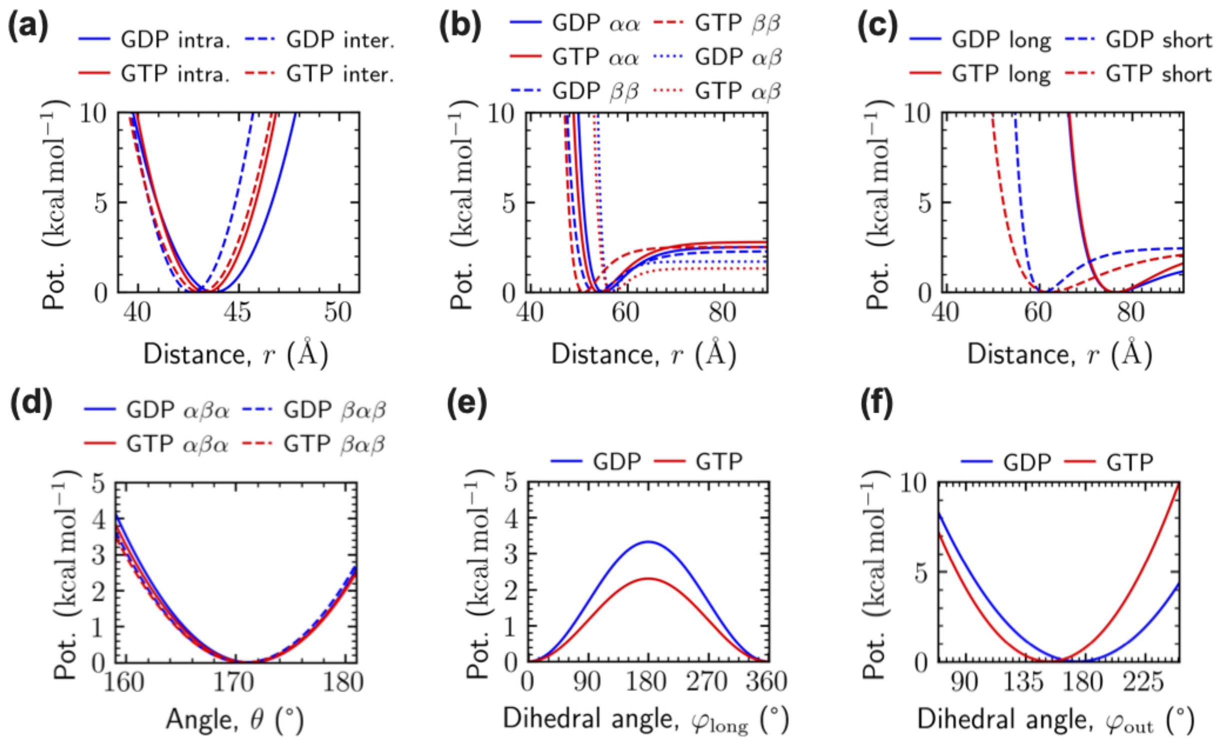


**FIGURE S5.** Comparison of analytic potentials parametrizing GDP- and GTP-MT. The analytic potentials are fits to the tabular potentials used in the IBI procedure. (a) intra- and interdimer distances; lateral (b) αα, ββ, and αβ (seam) lateral distances; (c) long and short diagonal distances; (d) αβα and βαβ longitudinal angles; (e) longitudinal dihedral angles, and (f) outward bias dihedral angles.

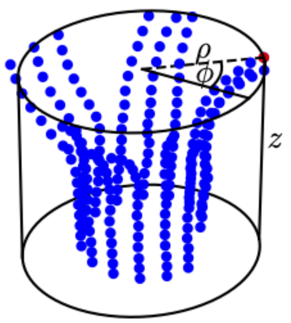


**FIGURE S6.** Cylindrical coordinate system used to analyze the MT plus-end tip trajectories.

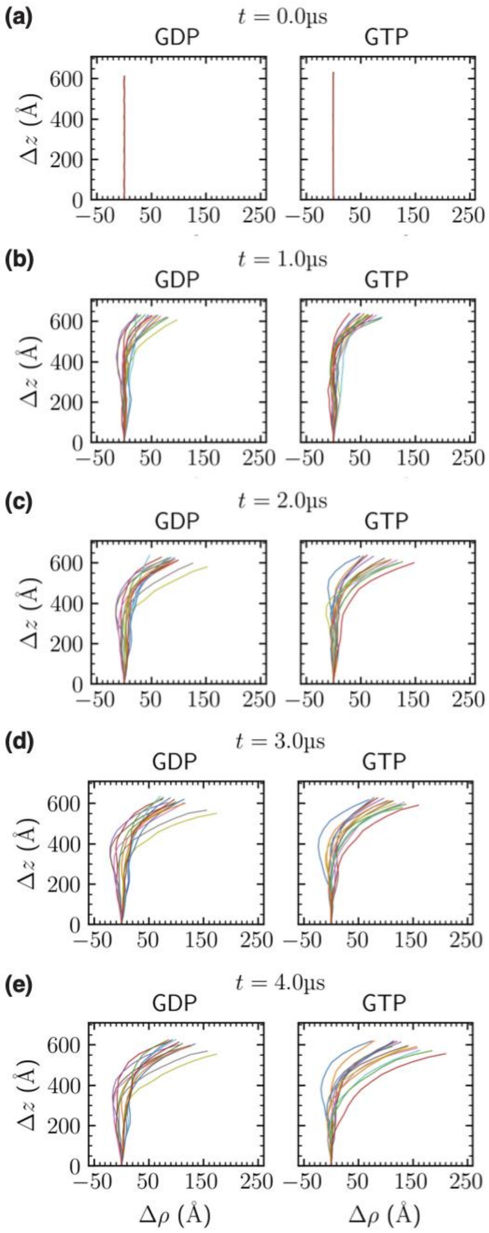


**FIGURE S7.** Radial slices presenting projections of PFs onto their individual radial planes at (a) 0, (b) 1, (c) 2, (d) 3, and (e) 4 µs.

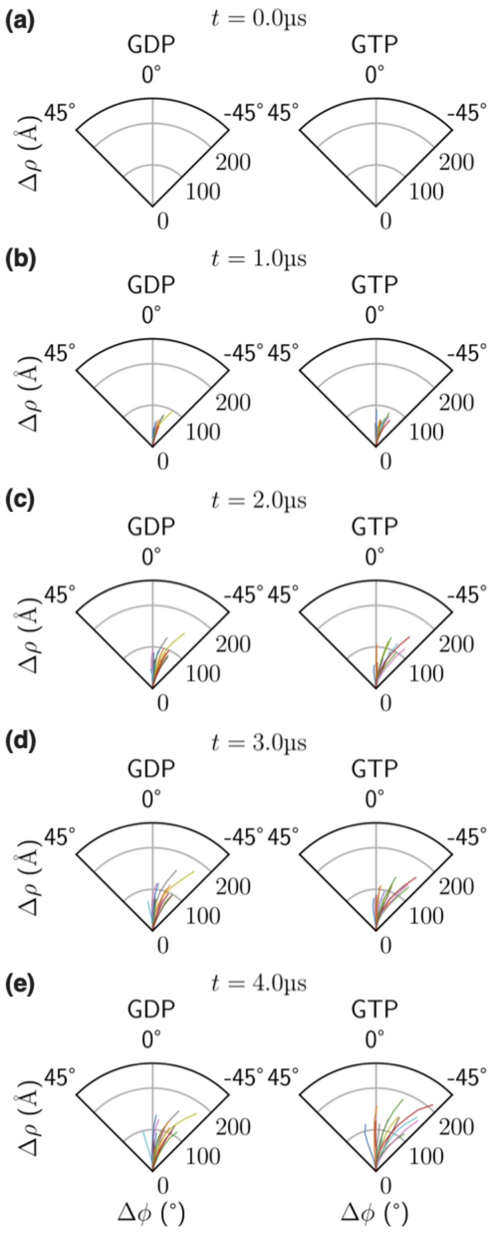


**FIGURE S8.** Axial slices presenting projections of PFs onto plane perpendicular to the MT axis. Polar plots show PFs at (a) 0, (b) 1, (c) 2, (d) 3, and (e) 4 μs.

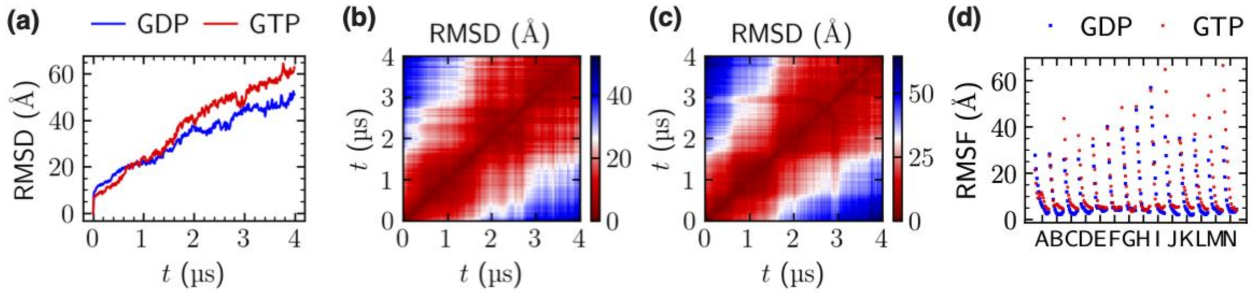


**FIGURE S9**. RMSD with respect to an initial, tubular form (a) and pairwise RMSD of (b) GDP- and (c) GTP-MT tips. (d) RMSF in monomer resolution from MD simulations.

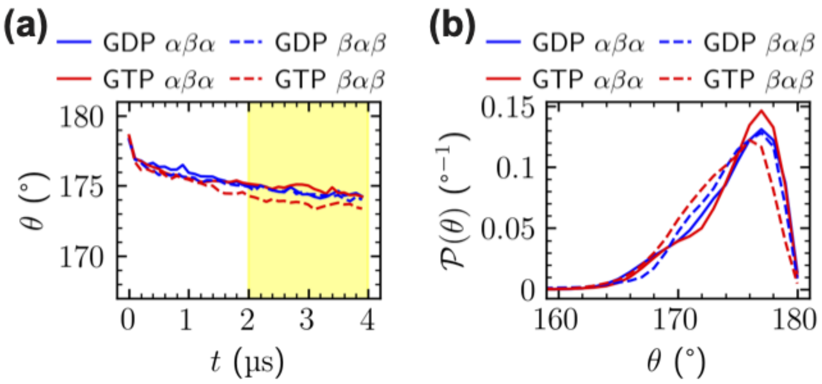


**FIGURE S10.** Longitudinal angles: (a) time evolution and (b) histograms of their values between 2 and 4 µs averaged over all symmetry-equivalent triples.

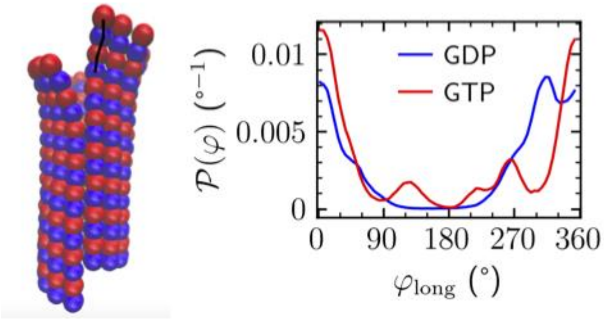


**FIGURE S11.** Longitudinal dihedral angles (example angle marked at the MT tip scheme): histogram of their values between 2 and 4 μs averaged over all topmost symmetry-equivalent quadruples.

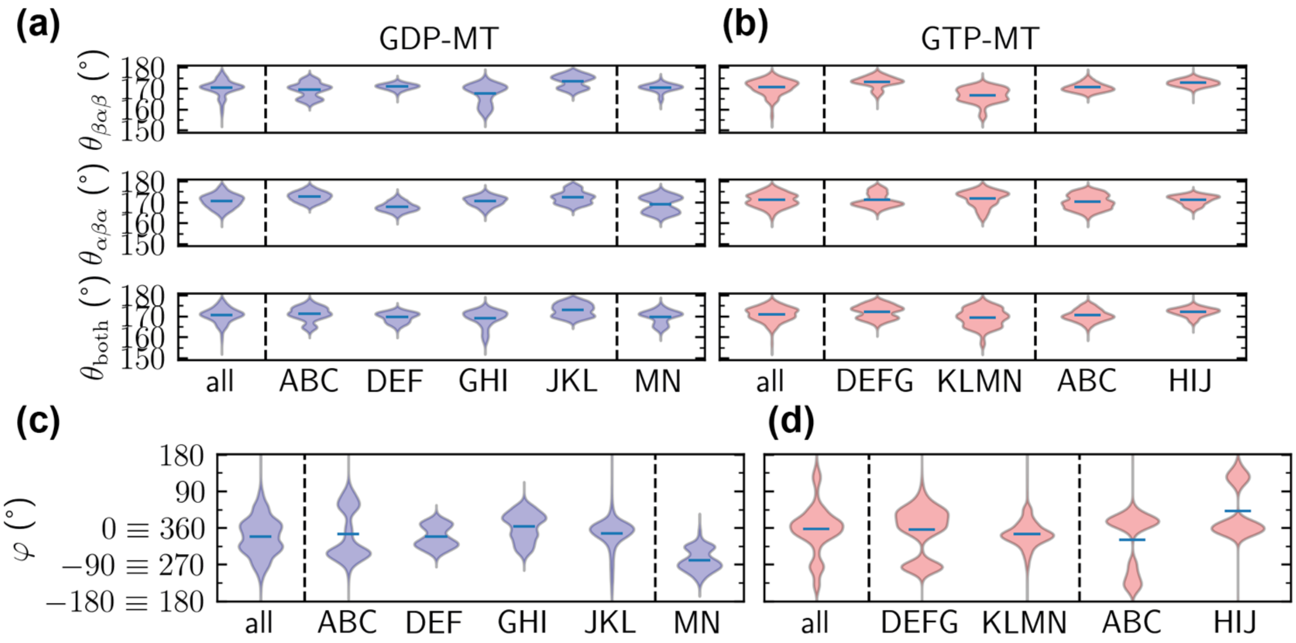


**FIGURE S12.** Histograms of longitudinal angles of tubulins belonging to specific protofilament clusters in (a) GDP- and (b) GTP-MT. Histograms of longitudinal dihedral angles of tubulins belongig to specific protofilament clusters in (c) GDP- and (d) GTP-MT.

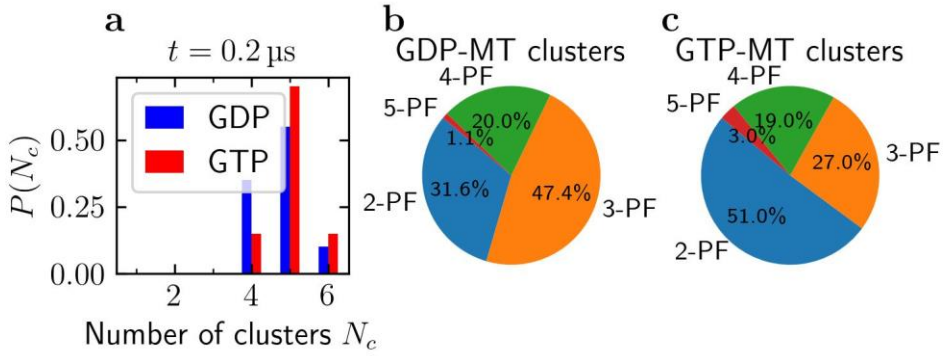


**FIGURE S13**. (a) Probability distribution of number of clusters computed by performing 20 0.2- μs-long independent Brownian dynamics simulations for GDP- and GTP-MT starting from microtubule tubula form. Percentages of clusters of given sizes in (b) GDP- and (c) GTP-MT.

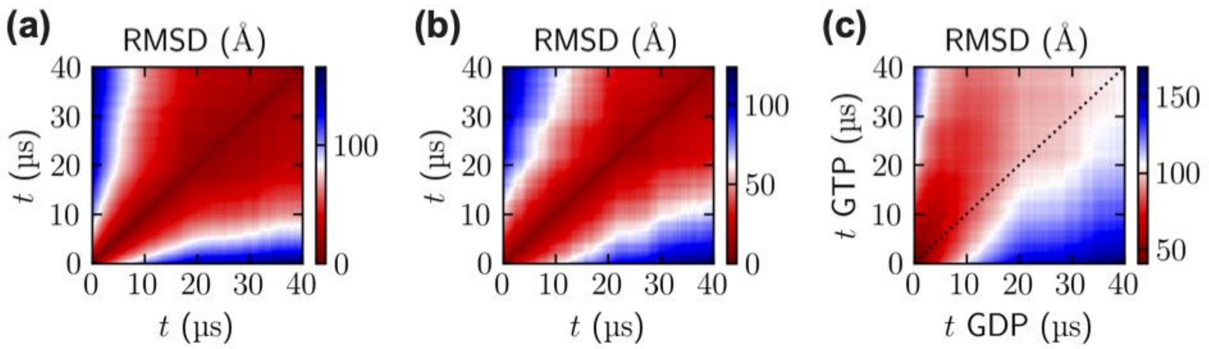


**FIGURE S14.** Pairwise RMSD of (a) GDP-MT, (b) GTP-MT, and (c) GDP- vs. GTP-MT.

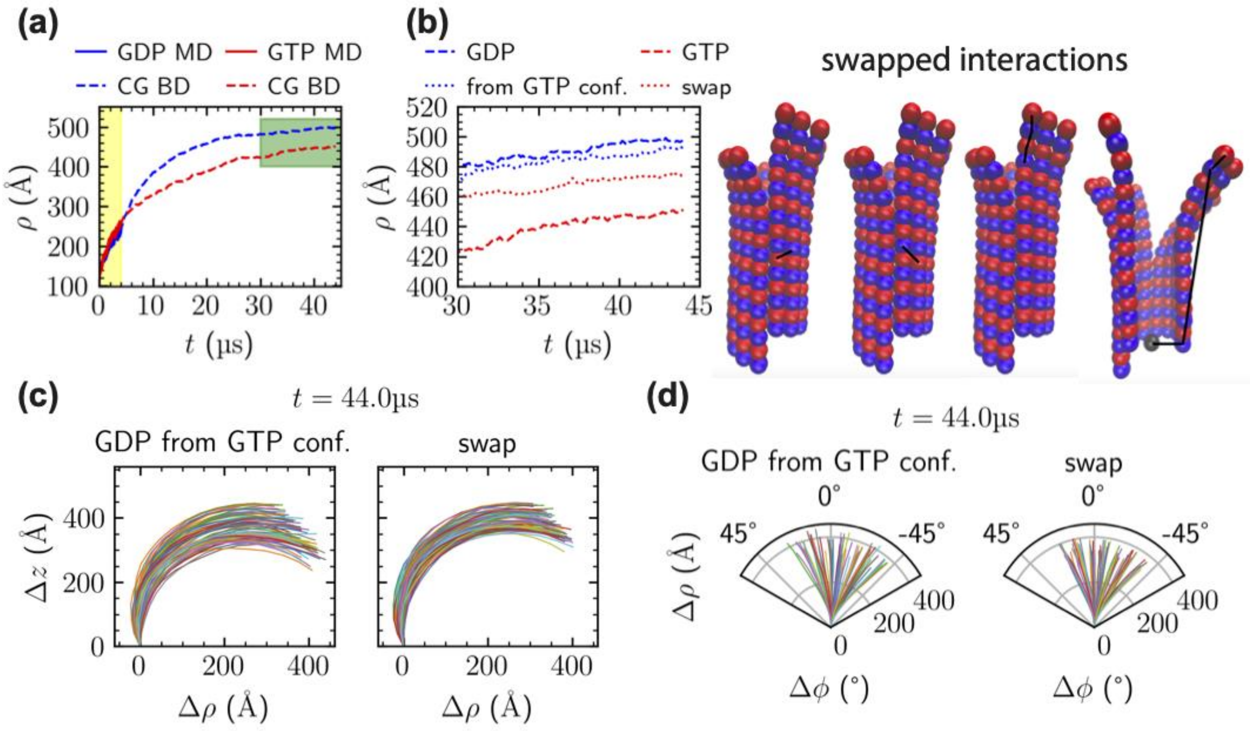


**FIGURE S15.** (a) Time evolution of radial distance ρ averaged over the topmost MT layer, and (b) a zoomed view on the 30-45 µs range. To determine if the observed differences between GDP- and GTP-MT tips were influenced by initial cluster composition, the following control simulations were performed: (i) GDP-MT plus-end tip model starting from initial configuration of GTP-MT ("GDP from GTP conf."), and (ii) GTP-MT plus-end tip model with 4 potentials exchanged for ones belonging to GDP-MT: 2 dihedral potentials, and 2 diagonal potentials ("swapped"). The first control did not substantially alter the long-term radial distance or PF shape. The second control reduced the difference in the average tip radial distance by half. (c) Projection of protofilaments onto their individual radial planes at 44µs for "GDP from GTP conf." and "swapped" systems. (d) Polar plots of protofilaments at 44µs for "GDP from GTP conf." and "swapped" systems.

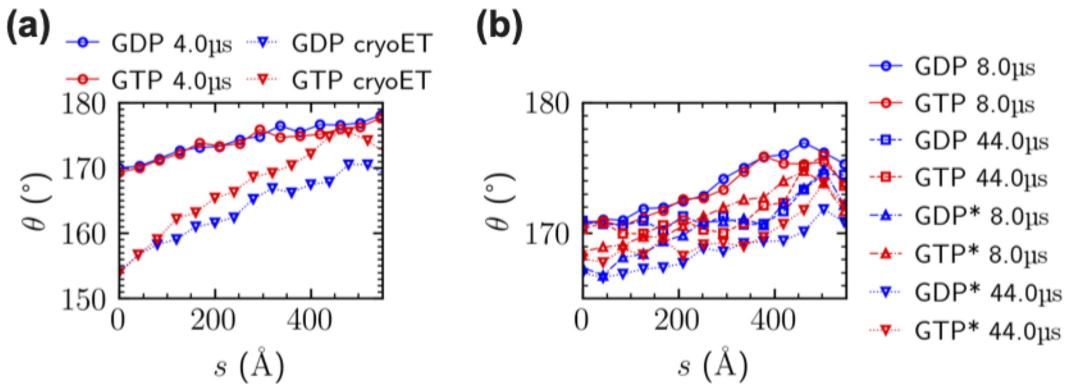


**FIGURE S16.** (a) Dependence of absolute curvature of the protofilaments on the distance from the tips from MD simulations and cryo-ET; (b) from CG BD simulations with and without top-down corrected angular potentials.

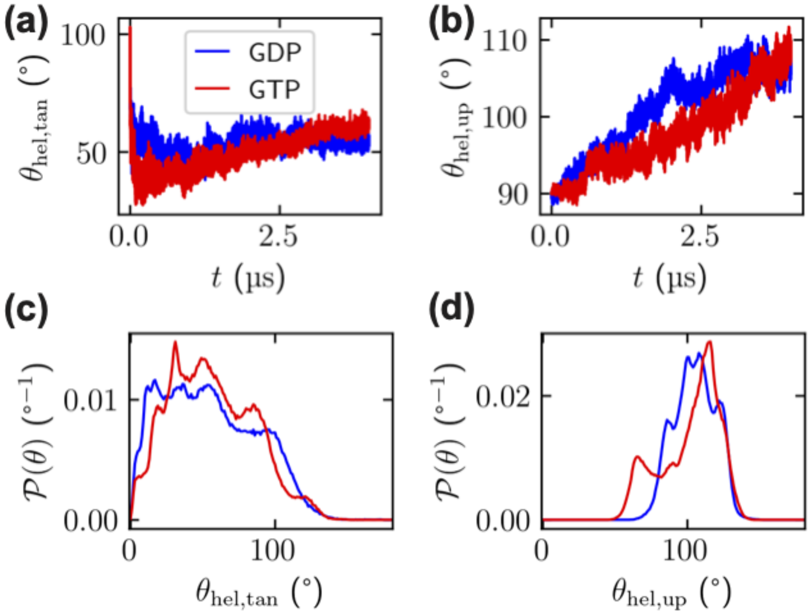


**FIGURE S17.** Orientation of vectors normal to PF plane: (a) time evolution of angle between the normal and counterclockwise tangential direction averaged over all topmost symmetry-equivalent triples; (b) time evolution of angle between the normal and MT axis averaged over all topmost symmetry-equivalent triples; (c,d) histograms of individual values which were averaged in (a,b)

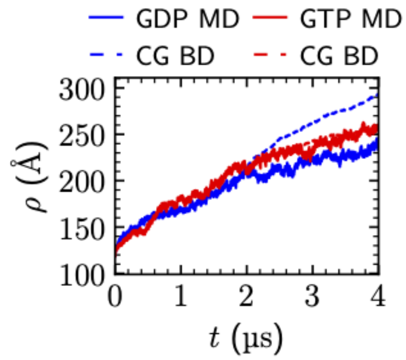


.

**FIGURE S18.** Radial coordinate of the CG MT tip models' topmost layers between 2 and 4 µs, compared to the dynamics from all-atom MD simulation.

**TABLE S1.** Details of box resizing in all-atom MD simulations of GDP- and GTP-MT tips.

| GDP | Time (μs) | MT protein atoms | Total atoms | $L_x$ (Å) | $L_y$ (Å) | $L_z$ (Å) |
|---|---|---|---|---|---|---|
| Stage 1 | 0 – 1.7 | 1,513,680 | 20,483,435 | 500 | 500 | 830 |
| Stage 2 | 1.7 - 3.9 | | 31,356,566 | 600 | 600 | 830 |
| Stage 3 | 3.9 - 4 | | 38,337,246 | 650 | 650 | 850 |
| GTP | | | | | | |
| Stage 1 | 0 – 1.5 | 1,514,240 | 20,981,887 | 500 | 500 | 850 |
| Stage 2 | 1.5 - 3.5 | | 32,118,650 | 600 | 600 | 850 |
| Stage 3 | 3.5 - 4 | | 39,232,415 | 650 | 650 | 870 |

**TABLE S2.** Statistical analysis of stable geometric characteristics of GDP- and GTP-MT plus-end tips from all-atom MD simulations. Values diferring statistically significantly for α=0.001 are bold. Values not differing significantly for α=0.1 are italic.

| | **Averaging** | **GDP mean** | **GTP mean** | **GDP std. dev.** | **GTP std. dev.** | **GDP eff. sample size** | **GTP eff. sample size** | **p value** |
|---|---|---|---|---|---|---|---|---|
| **Longitudinal intradimer distance** | all | **43.13** Å | **43.30** Å | 0.04 Å | 0.06 Å | 19 | 7 | $\mathbf{1.2\cdot10^{-4}}$ |
| | top | **43.27** Å | **43.81** Å | 0.08 Å | 0.14 Å | 73 | 12 | $\mathbf{1.2\cdot10^{-8}}$ |
| | bottom | **43.09** Å | **42.79** Å | 0.06 Å | 0.07 Å | 22 | 17 | $\mathbf{2\cdot10^{-15}}$ |
| **Longitudinal interdime distance** | all | **42.34** Å | **42.74** Å | 0.06 Å | 0.05 Å | 8 | 12 | $\mathbf{2\cdot10^{-10}}$ |
| | top | 42.7 Å | 42.89 Å | 0.2 Å | 0.12 Å | 6 | 47 | 0.08 |
| | bottom | 42.14 Å | 42.21 Å | 0.08 Å | 0.11 Å | 81 | 15 | 0.02 |
| **αβα longitudinal angle** | all | *174.5°* | *174.8°* | 0.2° | 0.3° | 6 | 6 | *0.14* |
| | top | 170.6° | 171.1° | 0.5° | 0.6° | 35 | 16 | 0.01 |
| **βαβ longitudinal angle** | all | **174.4°** | **173.7°** | 0.3° | 0.2° | 6 | 8 | $\mathbf{5\cdot10^{--4}}$ |
| | top | 170.3° | 170.7° | 0.5° | 0.6° | 49 | 16 | 0.07 |
| **Outwards-bias dihedral angle** | top | 155° | 148° | 2° | 3° | 15 | 5 | $5\cdot10^{-3}$ |

**TABLE S3.** Statistical analysis of dynamically changing geometric characteristics of GDP- and GTP-MT plus-end tips from all-atom MD simulations. Values diferring statistically significantly for α=0.001 are bold. Values not differing significantly for α=0.1 are italic.

| | Averaging | GDP trend | GDP intercept | GTP trend | GTP intercept | Error autocorrelation lag GDP | Error autocorrela[illegible] lag GTP |
|---|---|---|---|---|---|---|---|
| **αα lateral distance** | all | 2.04±0.08 Å/µs | *56.2±0.2* Å | 2.36±0.13 Å/µs | *56.6±0.4* Å | 498 | 745 |
| | top | *5.7±0.2* Å/µs | *62.6±0.7* Å | *6.1±0.3* Å/µs | *63.9±1.1* Å | 517 | 752 |
| | bottom | *0.048±0.009* Å/µs | *53.96±0.03* Å | *0.04±0.04* Å/µs | *53.97±0.13* Å | 181 | 1249 |
| **ββ lateral distance** | all | 2.20±0.08 Å/µs | *57.8±0.3* Å | 2.58±0.14 Å/µs | *58.2±0.5* Å | 508 | 745 |
| | top | *6.4±0.3* Å/µs | *70.2±0.8* Å | *7.0±0.4* Å/µs | *71.4±1.3* Å | 541 | 754 |
| | bottom | 0.029±0.004 Å/µs | 53.871±0.013 Å | 0.10±0.02 Å/µs | 53.69±0.08 Å | 70 | 865 |
| **Seam αβ lateral distance** | all | **0.8±0.2** Å/µs | *69.1±0.5* Å | **16.0±1.2** Å/µs | *62±4* Å | 148 | 1297 |
| | top | **5.7±0.9** Å/µs | **121±3** Å | **31±2** Å/µs | **151±7** Å | 162 | 722 |
| | bottom | **-0.85±0.08** Å/µs | **59.3±0.2** Å | **0.70±0.12** Å/µs | **55.4±0.4** Å | 251 | 338 |
| **Long diagonal distance** | all | **1.83±0.06** Å/µs | *77.4±0.2* Å | **2.97±0.09** Å/µs | *77.3±0.3* Å | 390 | 557 |
| | top | 6.7±0.3 Å/µs | *89.1±0.9* Å | 7.8±0.3 Å/µs | *88.5±1.1* Å | 601 | 710 |
| | bottom | *0.024±0.015* Å/µs | *74.47±0.04* Å | *0.04±0.02* Å/µs | *74.42±0.05* Å | 306 | 324 |
| **Short diagonal distance** | all | **1.60±0.07** Å/µs | *64.3±0.2* Å | **2.83±0.09** Å/µs | *64.5±0.3* Å | 424 | 567 |
| | top | *6.0±0.2* Å/µs | 72.3±0.8 Å | *6.0±0.4* Å/µs | 77.5±1.4 Å | 476 | 833 |
| | bottom | *-0.018±0.010* Å/µs | **62,34±0.03** Å | *-0.035±0.012* Å/µs | **61.99±0.04** Å | 147 | 185 |

TABLE S4. p-values from Welch t-test comparing means of αβα angle distributions across clusters and whole GDP-MT tip..

| | all | ABC | DEF | GHI | JKL | MN |
|---|---|---|---|---|---|---|
| all | | | | | | |
| ABC | 0 | | | | | |
| DEF | 0 | 0 | | | | |
| GHI | **0.72** | 0 | 0 | | | |
| JKL | $1.2\cdot10^{-14}$ | **0.08** | 0 | $1.4\cdot10^{-11}$ | | |
| MN | 0.00013 | $4\cdot10^{-15}$ | 0.007 | 0.0005 | $4\cdot10^{--12}$ | |

TABLE S5. p-values from Welch t-test comparing means of βαβ angle distributions across clusters and whole GDP-MT tip.

| | all | ABC | DEF | GHI | JKL | MN |
|---|---|---|---|---|---|---|
| all | | | | | | |
| ABC | 0.014 | | | | | |
| DEF | $7\cdot10^{-6}$ | $7\cdot10^{-5}$ | | | | |
| GHI | $3\cdot10^{-10}$ | 0.0012 | $2\cdot10^{-14}$ | | | |
| JKL | 0.0 | 0.0 | 0.0 | 0.0 | | |
| MN | **0.99** | 0.02 | 0.005 | $3\cdot10^{--9}$ | 0.0 | |

TABLE S6. p-values from Welch t-test comparing means of both angle distributions across clusters and whole GDP-MT tip.

| | all | ABC | DEF | GHI | JKL | MN |
|---|---|---|---|---|---|---|
| all | | | | | | |
| ABC | 0.002 | | | | | |
| DEF | 0 | 0 | | | | |
| GHI | $1.1\cdot10^{--9}$ | $9\cdot10^{--13}$ | **0.11** | | | |
| JKL | 0 | $2\cdot10^{-16}$ | 0 | 0 | | |
| MN | 0.0007 | $1.1\cdot10^{-6}$ | **0.28** | **0.05** | 0 | |

TABLE S7. p-values from Welch t-test comparing means of αβα angle distributions across clusters and whole GTP-MT tip.

| | all | DEFG | KLMN | ABC | HIJ |
|---|---|---|---|---|---|
| all | | | | | |
| DEFG | **0.97** | | | | |
| KLMN | 0.012 | **0.11** | | | |
| ABC | 0.007 | **0.05** | $5\cdot10^{-5}$ | | |
| HIJ | **0.71** | **0.78** | **0.09** | 0.008 | |

TABLE S8. p-values from Welch t-test comparing means of βαβ angle distributions across clusters and whole GTP-MT tip.

| | all | DEFG | KLMN | ABC | HIJ |
|---|---|---|---|---|---|
| all | | | | | |
| DEFG | 0 | | | | |
| KLMN | $2\cdot10^{--8}$ | $1.0\cdot10^{-14}$ | | | |
| ABC | **0.47** | 0 | $6\cdot10^{-8}$ | | |
| HIJ | $2\cdot10^{-16}$ | **0.11** | $7\cdot10^{-14}$ | 0 | |

TABLE S9. p-values from Welch t-test comparing means of both angle distributions across clusters and whole GTP-MT tip.

| | all | DEFG | KLMN | ABC | HIJ |
|---|---|---|---|---|---|
| all | | | | | |
| DEFG | $3\cdot10^{-11}$ | | | | |
| KLMN | $8\cdot10^{-11}$ | 0 | | | |
| ABC | 0.007 | $3\cdot10^{-14}$ | $8\cdot10^{-5}$ | | |
| HIJ | $6\cdot10^{-9}$ | **0.54** | 0 | $4\cdot10^{-12}$ | |

TABLE S10. p-values from Welch t-test comparing means of longitudinal dihedral angle distributions across clusters and whole GDP-MT tip.

| | all | ABC | DEF | GHI | JKL | MN |
|---|---|---|---|---|---|---|
| all | | | | | | |
| ABC | 0.04 | | | | | |
| DEF | **0.62** | 0.03 | | | | |
| GHI | 0 | $3\cdot10^{-12}$ | 0 | | | |
| JKL | 0 | **0.34** | $9\cdot10^{-14}$ | 0 | | |
| MN | 0 | 0 | 0 | 0 | 0 | |

TABLE S11. p-values from Welch t-test comparing means of longitudinal dihedral angle distributions across clusters and whole GTP-MT tip.

| | all | DEFG | KLMN | ABC | HIJ |
|---|---|---|---|---|---|
| all | | | | | |
| DEFG | 0.0009 | | | | |
| KLMN | 0 | 0 | | | |
| ABC | 0 | 0 | 0 | | |
| HIJ | 0 | 0 | 0 | 0 | |
